# The gut bacteria of an invasive amphibian respond to the dual challenges of range-expansion and parasite attack

**DOI:** 10.1101/2020.11.16.385690

**Authors:** Jia Zhou, Tiffanie Maree Nelson, Carlos Rodriguez Lopez, Shao Jia Zhou, Georgia Ward-Fear, Katarina C. Stuart, Richard Shine, Lee Ann Rollins

**Author notes:** **Correspondence:** Jia Zhou.

## Abstract

Gut bacterial communities influence, and are influenced by, the behaviour and ecology of their hosts. Those interactions have been studied primarily in humans and model organisms, but we need field research to understand the relationship between an organism’s gut bacteria and its ecological challenges, such as those imposed by rapid range expansion (as in invasive species) and the presence of host-manipulating parasites. Cane toads (*Rhinella marina*) provide an excellent model system in this respect, because the species’ ongoing colonization of Australia has enforced major changes in phenotypic traits (including behaviour), and lungworm parasites (*Rhabdias pseudosphaerocephala*) modify host gut function in ways that enhance the viability of lungworm larvae. We collected female toads from across the species’ invasive range and studied their morphology, behaviour, parasite infection status and gut bacterial community. Range-core *versus* range-edge toads differed in morphology, behaviour, gut bacterial composition and predicted gut bacterial function but did not differ in the occurrence of parasite infection nor in the intensity of infection. Toads infected with lungworms differed from uninfected conspecifics in gut bacterial composition and diversity. Our study demonstrates strong associations between gut bacterial community and host ecology and behaviour.

## Introduction

The bacterial community within an organism’s intestines can be strongly influenced by host behaviour and ecology, such as habitat selection and diet [1–5]. But that interaction runs both ways because gut bacteria can influence behaviour and ecology of the host. For example, transferring gut contents can modify the recipient host’s behaviour (exploratory behaviour, *Mus musculus* [6]; emotional reactivity, *Coturnix japonica* [7]). Similarly, altering gut microbial communities by administering antibiotics or altering dietary composition triggered aggressive behaviour in leaf-cutting ants (*Acromyrmex echinatior* [8]). Remarkably, changes in only a single bacterial species within the gut can affect behaviour of the host (e.g., *Drosophila melanogaster* [9]; *Danio rerio* [10]; *M. musculus* [11]). Gut microbiota can also affect mating choices [12] and foraging [13–15].

To date, most evidence for effects of intestinal bacteria on host behaviour comes from studies on humans and “model organisms”. To elucidate the functional significance of this phenomenon, we need to extend such studies to free-ranging animals, incorporating a wider range of taxa [16]. Invasive species offer good models for such research, because novel challenges in the invaded range create an opportunity to compare closely-related organisms exposed to profoundly different environments [17,18]. Host-parasite relationships also may be disrupted during biological invasions, due to processes such as “enemy release” (loss of co-evolved native pathogens from the native range [19]). Parasites can manipulate host behaviour and physiology in ways that enhance parasite fitness but reduce host fitness [20]. Thus, if the gut bacterial community provides a mechanism for such effects, parasite-infected individuals should exhibit different gut bacteria than uninfected conspecifics.

These ideas suggest two predictions: (i) that the gut bacterial community should differ between populations of an invasive species (e.g., between the range-core and the invasion-front); and (ii) that the gut bacterial community should differ between parasitized hosts and non-parasitized conspecifics. The colonization of Australia by cane toads (*Rhinella marina*) provides a robust opportunity to test these predictions. Since their release in north-eastern Australia in 1935, toads have dispersed into areas that are much hotter and more seasonally arid than in the native range or the initial release sites [21]; and toads have brought with them a native-range nematode lungworm (*Rhabdias pseudosphaerocephala*) that can have devastating impacts on host viability, and induces behavioural and physiological changes in the host [22]. Notably, infected toads produce copious watery faeces [22]; hence, we expect lungworm-infected toads to exhibit different gut bacterial communities than uninfected individuals.

## Methodology

### Study species, sample collection and behavioural assays

Cane toads are native to South America, and were introduced into Australia in 1935 as a biocontrol for pests of sugar cane crops [23]. As the toads spread through tropical Australia, they fatally poisoned many native predators [23]. Toads from range-core populations (eastern Australia) differ from invasion-front conspecifics (in north-western Australia) in phenotypic traits that confer increased dispersal ability, such as endurance [24], limb morphology [25], boldness and exploratory behaviour [17,26]. Toads from the invasion-front also have lower rates of infection of the co-introduced parasitic lungworm [27]. Drivers of variation in invasion-related behaviours in this species include genetics, morphology, habitat, diet, prior experience and parasites [17,28–31]. However, the possible role of gut bacteria as a potential driver of behavioural shifts across the invasive range has not been studied.

We hand-captured 60 adult females from three sites on the invasion-front and three sites in the range-core (Table S1). We conducted brief behavioural assays upon collection including: (i) struggle score (number of kicks after being captured until toad remains still for 5 seconds) and struggle likelihood; and (ii) righting effort (time to right itself, number of kicks within two minutes after toad is placed on its dorsal side, and righting effort likelihood [17]. These measures are predictive of traits including speed and stamina (K. Stuart, pers. comm.), suggesting that these simple assays may reveal a toad’s dispersal potential. We then placed animals into individual, moist, calico bags and weighed, measured (snout urostyle length; SUL) and euthanised them by injecting tricaine methanesulfonate (MS222) buffered with bicarbonate of soda.

We dissected the toads and scored the presence of two types of toad parasite: the gut-encysted physalopterine larvae [32] and adult lungworms [22,33]. Lungworm larvae pass through the gut, but are less easily detected and reliably counted than are adult lungworms. From each toad, we removed 0.3cm of colon near the cloaca (including gut contents) and preserved it in 95% ethanol (see Supplementary Material for justification of sampling protocols).

### Analyses

We compared host morphology and behaviour between regions (range-core *versus* invasion-front), and as a function of infection status (lungworm infected *versus* non-infected). Because body length (SUL) and mass were correlated, we only included SUL as our measure of host morphology in further analyses. We used a t-test to compare mean SUL between regions and infected/non-infected toads. For associations between region or infection status with host characteristics or behavioural traits, we used SUL as a covariate in generalized linear models (GLM). See Supplementary Material for details of statistical analyses.

Laboratory methods and data pre-processing for characterizing gut bacterial community composition and predicted functions are described in Supplementary Material. Briefly, we calculated within-individual (alpha) bacterial diversity and between-site (beta) bacterial diversity. For the latter variable, we subset our data to include the Core50 gut community (Amplified Sequence Variants (ASVs) present in a minimum of 50% of toads from each site [2]). We predicted bacterial functions and generated pathway abundance based on Core50 ASVs. We compared bacterial composition and predicted function between regions, and between lungworm-infected and non-infected toads. We identified differences in individual ASVs and predicted bacterial functions between range-core and invasion-front toads and identified associations between host characteristics (including infection with parasites) with bacterial communities and predicted bacterial functions. Analyses were conducted in QIIME2 [34], PICRUST2 [35], and R packages in R version 4.0.2 [36].

## Results

### Ecological traits

Wild-caught invasion-front toads were larger than range-core toads (Tables S2, S3; mean SUL t = 2.54, df = 53.90, p = 0.014). Neither counts nor presence of parasites (lungworm and gut) differed significantly across the range (Table S2). Range-core toads were more likely to struggle (Tables S2, S3; p = 0.008, 95% CI: core [−0.174, 0.005], edge [−0.023, 0.199]) and, in those that did struggle, the number of struggle movements was higher for range-core toads (Tables S2, S3; p = 0.002, 95% CI: core [−0.057, 0.026], edge [0.046, 0.2]). Range-core toads also were more likely to attempt to right themselves (p = 0.036, 95% CI: core [−0.092, 0.04], edge [−0.006, 0.19]), but righting effort and righting time did not differ significantly between geographic regions (Tables S2, S3).

Because there were no significant differences in prevalence or intensity of lungworm between the range-core and invasion-front toads (Table S2), we combined samples to analyse correlates of lungworm infection. Infected toads were similar in SUL to non-infected toads (Tables S3, 4; t = 0.86, df = 57.19, p = 0.393), with no significant behavioural differences between the two groups (Table S3, 4).

### Gut bacterial community composition and predicted bacterial function

Alpha diversity did not differ significantly between regions (Supplementary Material), but beta diversity of bacterial taxonomic communities differed between regions (Figures 1A, S1; R^2^ = 0.050, F = 3.050, p < 0.001) and sampling sites (Table S5; all p-values < 0.001). Among 230 ASVs that were assigned to family level, the abundance of 124 ASVs differed between the colons of range-core *versus* invasion-front toads (Table S6). The number of significantly different ASVs in each phylum were: Bacteroidetes (60 ASVs), Firmicutes (55 ASVs), Proteobacteria (7 ASVs), Actinobacteria (1 ASVs), Verrucomicrobia (1 ASV) (Table S6, Figure 2A).

**Figure 1.**
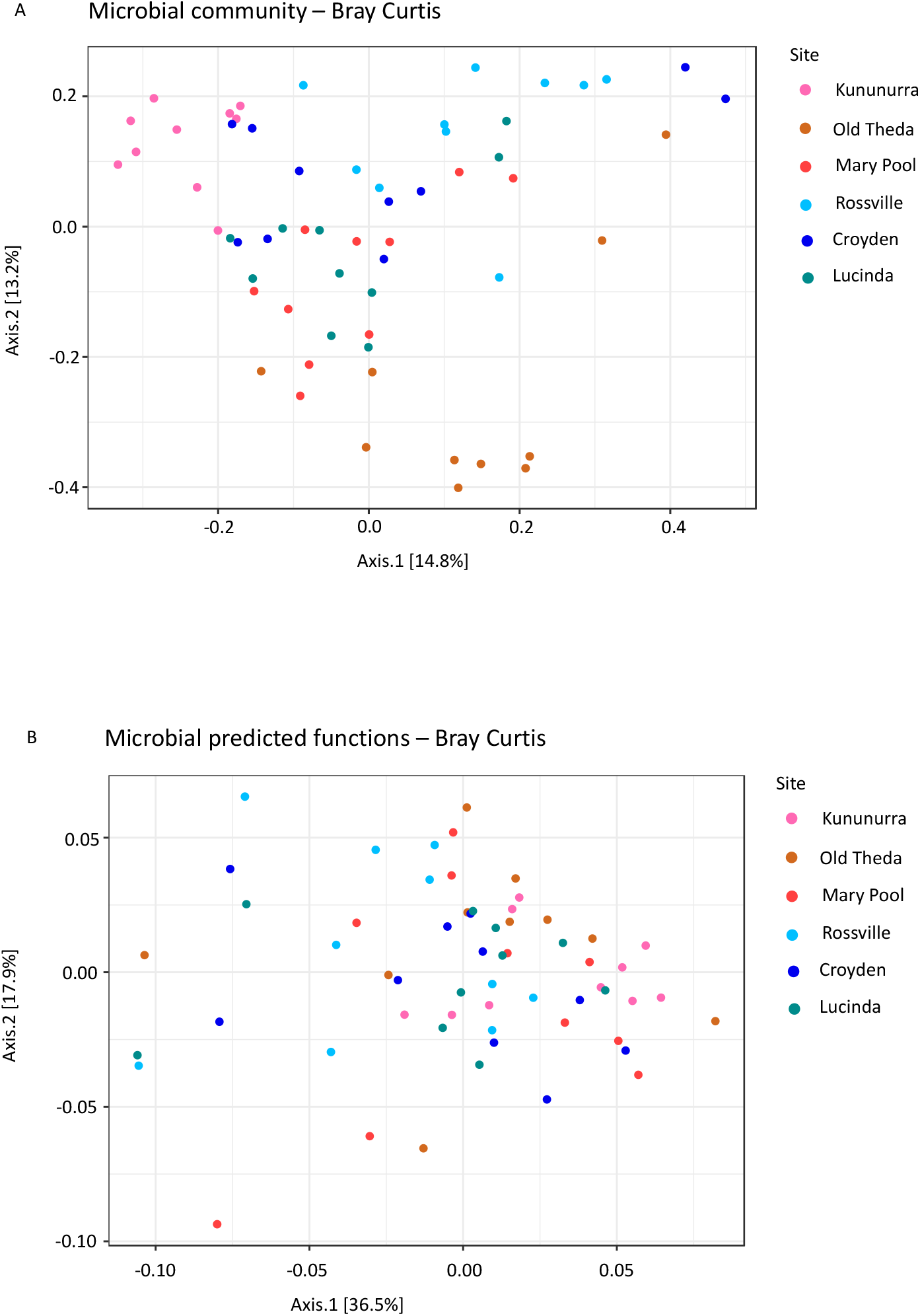
Beta diversity by location. Principle coordinate analysis plot of Bray Curtis distance of microbial community (A) and Bray Curtis distance of predicted functional groups (B) from 60 cane toad individuals of the invasion-front (Kununurra, Old Theda, and Mary Pool) and the range-core (Rossville, Croydon, and Lucinda).

**Figure 2.**
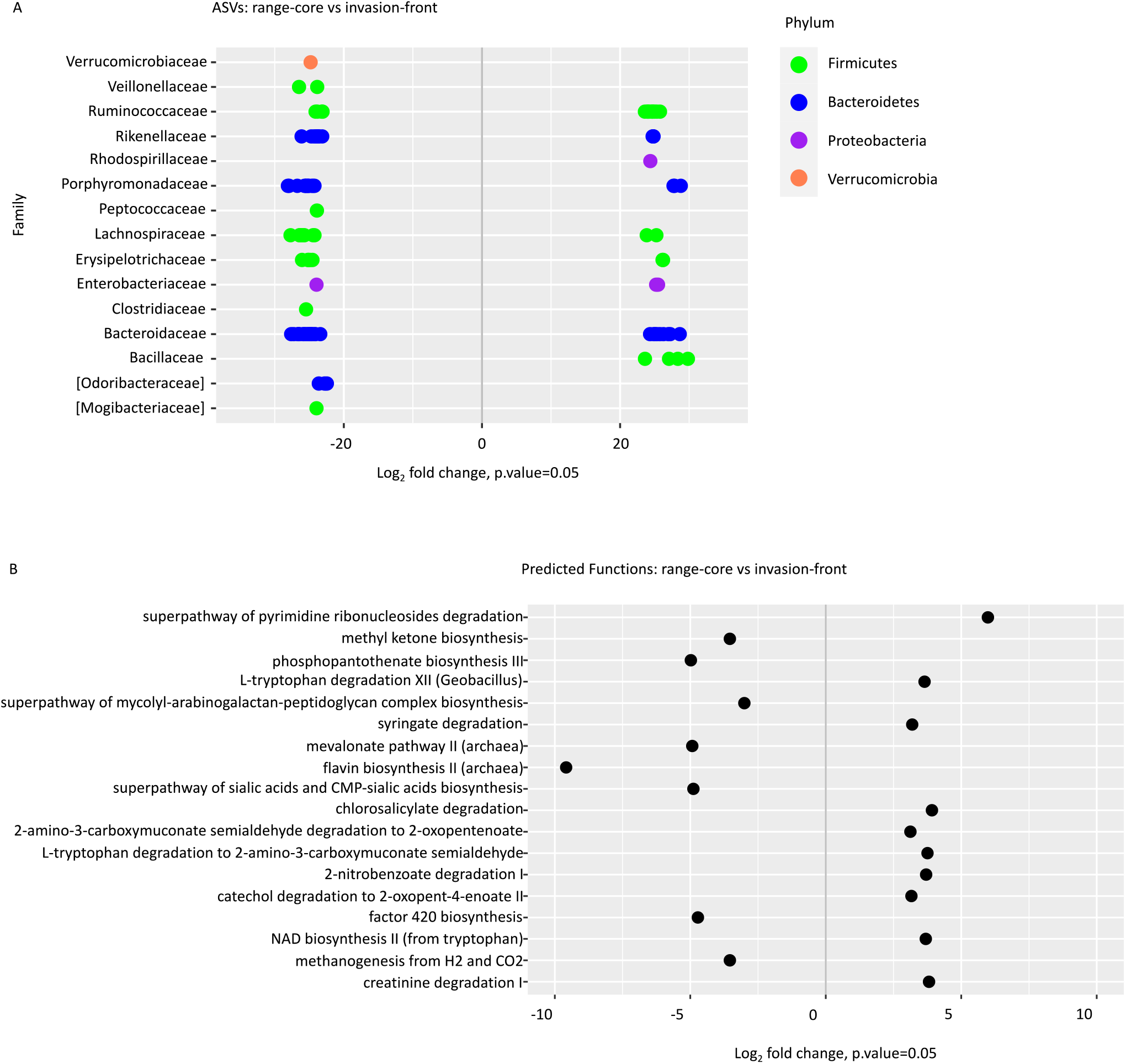
Significantly different bacterial taxa and predicted functions between range-core (QLD) and invasion-front (WA) toads’ colon. Significant differences were identified between locations via differential abundance testing based on a negative binomial distribution. The dots represent the average log-2 fold change (x axis) abundance and positive log2 fold changes signify increased abundance in range-core, and negative log_2_ fold changes display increased abundance in invasion-front. Bacterial taxa (A) were classified to the taxonomic level of family (y axis) and coloured by taxonomic level of phylum. Family name in bracket is proposed taxonomy by Greengenes. Only ASVs that could be matched to a known bacterial family and with a log2FoldChange value higher than 20 or lower than −20 are presented. Predicted functions (B) with a log2FoldChange value higher than 3 or lower than −3 are presented.

Among the identified 474 predicted bacterial functions, we found significant differences between invasion-front and range-core toads (Figure 1B; R^2^ = 0.064, F = 4.110, p-value= 0.002). Pairwise tests between sampling sites indicated that Kununurra toads had different bacterial functions to Rossville (p-value= 0.009) and Lucinda toads (p-value = 0.046), but no other sites differed functionally from each other (Table S7; all other p-values > 0.05). In total, 84 predicted bacterial functions differed between invasion-front and range-core toads (Table S8, Figure 2B). Range-core toads had more abundant bacterial function in the superpathway of pyrimidine ribonucleosides degradation (log2FoldChange = 5.98) and less abundant bacterial function in phosphopantothenate biosynthesis III (log2FoldChange = −4.98), superpathway of sialic acids and CMP-sialic acids biosynthesis (log2FoldChange = −4.89) and factor 420 biosynthesis (log2FoldChange = −4.72) than did invasion-front toads (Table S8, Figure 2B). Among the 30 most abundant functions, range-core toads had lower bacterial function in urate biosynthesis/inosine 5’-phosphate degradation (log2FoldChange = −0.10) than did invasion-front toads (Figure 3, Table S8).

**Figure 3.**
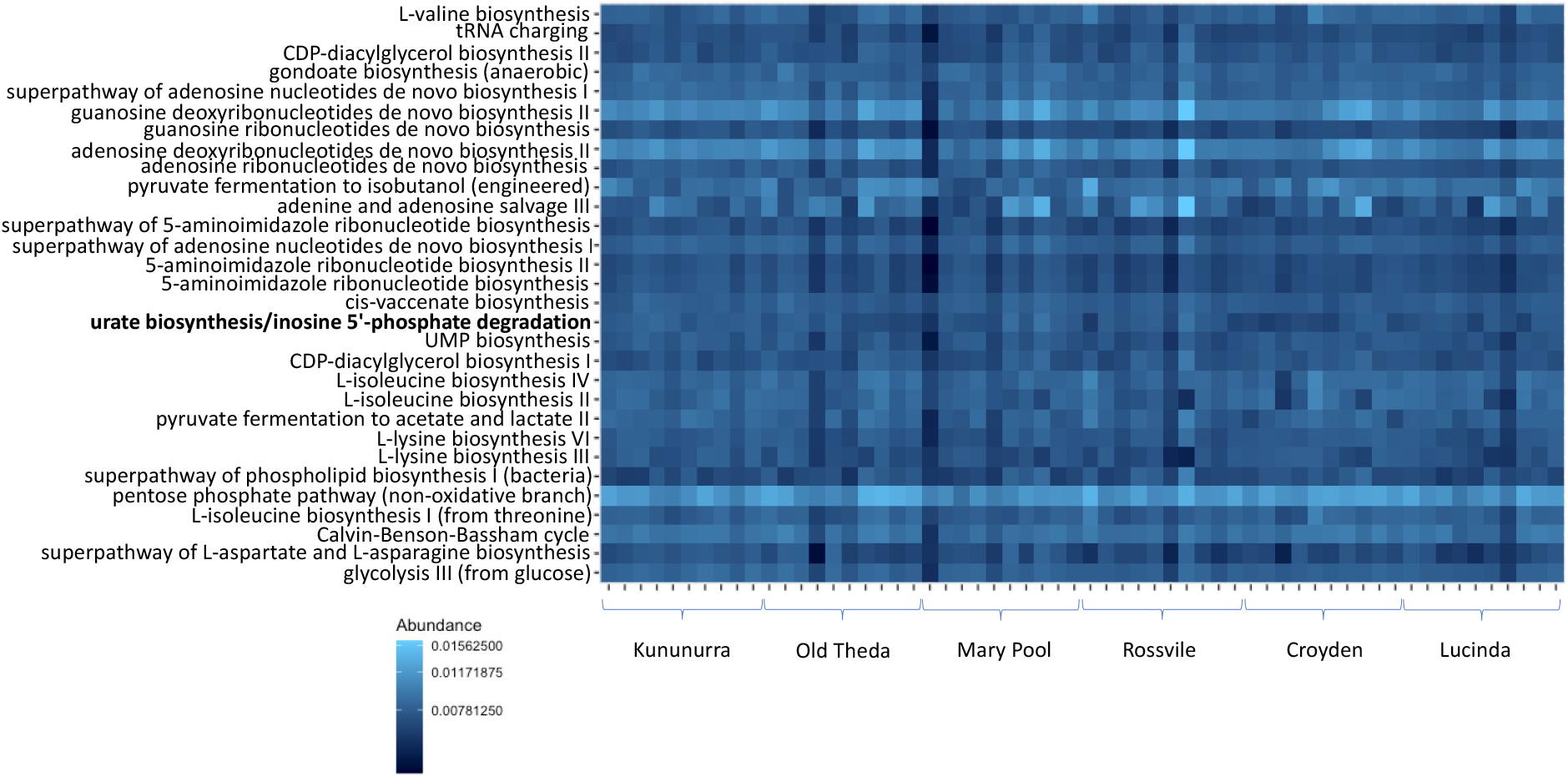
Heatmap for top 30 functional group abundance. Heatmap indicates the top 30 functional groups in the intestinal samples from range-core and invasion-front toads. Abundance indicates the raw count of functional groups inferred from taxonomic 16S sequences using PICRUSt where light blue is high abundance and dark blue is lower abundance. Functional pathways that differ significantly between range-core and invasion-front toads are highlighted in bold. Range-core includes Rossville, Croyden, and Lucinda; invasion-front includes Kununurra, Old Theda, and Mary Pool.

### Associations between ecological traits and intestinal bacteria

To assess correlates of gut bacterial composition and function, we compared characteristics of individual hosts to bacterial variation. Only the occurrence of lungworms was significantly associated with the bacterial composition (R^2^ = 0.128, p = 0.02) (Table 1; Figure S2A, B).

**Table 1a-d.** The association between a) single host factors and microbial community, b) single host factors and predicted microbial function, c) single behavioural trait and microbial community and d) single behavioural trait and predicted microbial function. Significant p-values denoted by an asterisk.

|  | MDS1 | MDS2 | r<sup>2</sup> | Pr(>r) |
| --- | --- | --- | --- | --- |
| *a. Host factor/microbial community* |  |  |  |  |
| SUL | 0.170 | 0.985 | 0.071 | 0.119 |
| BodyWeight | -0.009 | 1.000 | 0.051 | 0.231 |
| Lungworms | 0.254 | -0.967 | 0.020 | 0.585 |
| Occurrence of lungworms | 0.453 | -0.892 | 0.128 | 0.023* |
| *b. Host factor/microbial function* |  |  |  |  |
| SUL | 0.680 | 0.733 | 0.021 | 0.556 |
| BodyWeight | 0.610 | 0.793 | 0.041 | 0.311 |
| Lungworms | -0.791 | 0.612 | 0.007 | 0.839 |
| Occurrence of lungworms | -0.827 | -0.562 | 0.059 | 0.187 |
| *c. Behavioural trait/microbial community* |  |  |  |  |
| Struggle score | -0.814 | 0.580 | 0.044 | 0.268 |
| Struggle likelihood | -0.880 | -0.474 | 0.021 | 0.555 |
| Righting effort | -0.124 | 0.992 | 0.075 | 0.121 |
| Righting effort likelihood | 0.499 | 0.867 | 0.0255 | 0.474 |
| Righting time | 0.933 | 0.360 | 0.0615 | 0.174 |
| *d. Behavioural trait/microbial function* |  |  |  |  |
| Struggle score | 0.376 | 0.927 | 0.051 | 0.221 |
| Struggle likelihood | 0.183 | 0.983 | 0.017 | 0.606 |
| Righting effort | 0.571 | 0.821 | 0.070 | 0.141 |
| Righting effort likelihood | 0.399 | 0.917 | 0.035 | 0.362 |
| Righting time | -0.956 | -0.294 | 0.004 | 0.894 |

In a redundancy analysis combining ecological traits measured here, the model that explained the most variation in the bacterial community assemblage included only the occurrence of lungworms (AIC = 178.58). The best model to explain variation in predicted bacterial functions included the likelihood of righting (AIC = 53.613), the occurrence of lungworms (AIC = 54.297) and righting time (AIC = 56.912). The combination of these three factors explained 17.8% of total variation in predicted bacterial functions (Figure 4).

**Figure 4.**
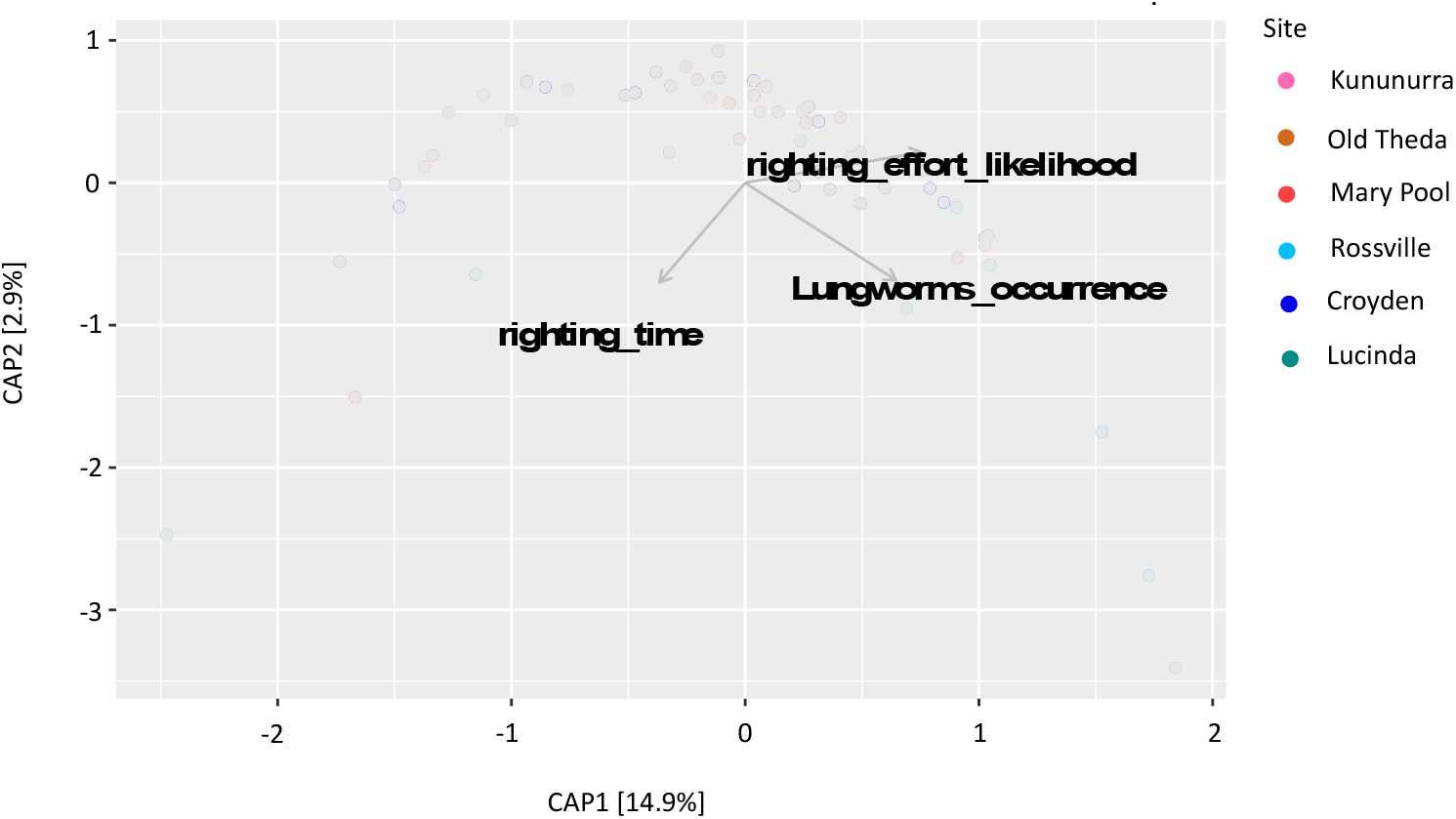
Main variables that affect predicted function differentiation among individuals. CAP (*capscale*) plot displays the combination of variables that explained the greatest variation in the predicted functions through model selection, using 60 cane toad individuals from the invasion-front (Kununurra, Old Theda, and Mary Pool) and the range-core (Rossville, Croydon, and Lucinda). The final model explained 17.8% of variation in the microbial predicted functions, which includes righting effort likelihood (AIC = 53.613), occurrence of lung worms (AIC = 54.297) and righting time (AIC = 56.912) explained the greatest variation.

In explicit tests of whether bacterial community assemblages differed in infected *versus* non-infected toads, we found a significant association with lungworm occurrence (Table 2, p=0.005). Intensity of lungworm infection was not significantly associated with gut bacterial community but did have a significant interaction with region in this analysis (Table 2, p=0.04).

**Table 2.** Association between gut microbiota variation and lungworm occurrence and intensity, based on Bray Curtis dissimilarity values for microbial community assemblages.

|  | Df | SumOfSqs | R2 | F | Pr(>F) |
| --- | --- | --- | --- | --- | --- |
| Location | 1 | 1.314 | 0.091 | 6.026 | <0.001*** |
| Lungworm_occurrence | 1 | 0.482 | 0.033 | 2.213 | 0.005** |
| Lungworms_intensity | 1 | 0.150 | 0.010 | 0.688 | 0.861 |
| Location:Lungworm_occurrence | 1 | 0.331 | 0.023 | 1.520 | 0.075 |
| Location:Lungworm_intensity | 1 | 0.357 | 0.025 | 1.638 | 0.043* |
| Residual | 54 | 11.774 | 0.817 |  |  |
| Total | 59 | 14.409 | 1.000 |  |  |

Significance codes: 0 '\*\*\*' ≤0.001 '\*\*' ≤0.01 '\*' ≤0.05.

## Discussion

Bacteria influence animal behaviour in diverse ways [16,37], but the ecological drivers of variation in gut bacterial composition remain largely unstudied. Our analyses of cane toads from two regions within their invasive range documents substantial variation in community assemblage and function of gut bacteria. Importantly, that variation was associated with two traits that we predicted to influence gut bacterial assemblages: invasion history and parasite infection. Interestingly, toad behaviour differed across the invasive range, and toad righting behaviour was associated with bacterial function but not with parasite infection.

### Geographic divergence in gut bacteria

First, we consider the differences in gut bacteria between toads from the invasion-front and the range-core. Although these populations have been separated by less than a century, the toads have diverged remarkably in morphology, physiology and behaviour and much of that divergence is heritable [38,39]. Some of those shifts likely reflect evolutionary pressures for increased rates of dispersal, due to adaptive (natural selection) and non-adaptive (spatial sorting) mechanisms [40,41]. Other geographically variable aspects of toad phenotypes likely are responses to different climatic conditions in the newly-invaded regions (hot, seasonally arid) compared to the range-core (cooler, more equable climate) [42]. Similar geographic divergence has been reported for the microbiome on the toad’s skin [43]. Our data illustrate that the invasion of Australia by cane toads has been accompanied by substantial divergence in gut bacterial communities. Alpha diversity in gut bacteria was similar in invasion-front and range-core individuals, but there were differences in both the gut bacteria composition and predicted bacterial function between toad populations across the species’ Australian invasive range. Predicted bacterial functions better explained cane toad righting behaviour than did gut bacterial community composition. Intriguingly, similarity between gut bacterial communities between individuals within regions in Australia is related to the similarity of their host’s epigenome, and this relationship is strengthened in populations where genetic diversity is lowest, such as on the invasion front [44]. Relationships between gut bacterial communities and their hosts are complex, and that a clear understanding of these relationships requires careful consideration of numerous environmental, host and gut bacterial factors.

The diversity and composition of bacterial communities differed between range-core and invasion-front toads, despite an overall similarity in their dominant phyla and alpha diversity. ASVs in the family Veillonellaceae were higher at the invasion-front (Figure 2A). The abundance of this bacterial family may influence host metabolic regulation. For example, in Brandt’s voles (*Lasiopodomys brandtii*) exposed to colder temperatures, voles which huddled had more Veillonellaceae and more short-chain fatty acids (SCFAs) in their intestines than did non-huddling voles [45]. This family produces SCFAs such as propionic acid [46,47], which can increase locomotor activity [48]. The link to host metabolic regulation suggests that invasion-front toads might fuel their invasion in this way [24]. ASVs from another family of SCFA-producing bacteria, Clostridiaceae [49], were also higher in invasion-front toads than those from the range-core. Furthermore, the family Veillonellaceae may be associated with host sociality. A reduction of Veillonellaceae has been observed in children with Autism Spectrum Disorder, often known for desiring social isolation [50]. Higher abundance of Veillonellaceae in invasion-front toads could foster their “bolder” personality, retaining a higher propensity for exploration and risk-taking [26,51].

Several other ASVs that differed across the toad’s range also may affect behaviour. ASVs from the family Peptococcaceae, more common in invasion-front toads (Figure 2A), are related to host neurotransmitter levels (noradrenaline linking visual awareness to external world events [52]). For example, Peptococcaceae levels in the caecum are positively correlated with noradrenaline levels in mice [53]. ASVs from family Bacillaceae, lower in invasion-front toads (Figure 2A), might be related to host anxiety (e.g., abundant in methamphetamine-treated rats [54], and in exercised *versus* sedentary mice [55]). Abundant Bacillaceae might induce anxiety-like behaviours, thus intensifying the stress response [54] and decreasing exploratory behaviour in new environments [56]. In summary, invasion-front toads possessed gut bacterial communities that in other studies have been associated with SCFAs production and neurotransmitters. That pattern supports the idea that gut microbes in invasion-front toads may increase locomotor ability, alertness and propensity for exploration and risk-taking. In comparison, range-core toads possessed bacterial taxa that have been associated with anxiety, and a decreased propensity to explore.

Geographic variation was less obvious in the predicted bacterial functional groups than in community composition (Figure 1), consistent with the hypothesis that bacterial function is more conservative than taxonomic composition (e.g. in fire salamanders [2]). Different gut microbiota can have similar bacterial functions, increasing resilience and functional stability [2,3,57]. Despite this broad similarity, bacterial functions differed between range-core and invasion-front toads. These differences included those involved in functional pathways related to food sources and metabolism. Invasion-front toads had less bacterial function in the superpathway of pyrimidine ribonucleosides degradation, which provides a nitrogen source for microbes [58] and plays an important role in perturbations in the uridine monophosphate (UMP) biosynthetic pathways. This pathway allows the bacterial cell to sense signals such as starvation, nucleic acid degradation, and availability of exogenous pyrimidines, and to adapt the production of the extracellular matrix to changing environmental conditions [59]. This function might help to explain the disappearance of Verrucomicrobia as a dominant taxon. As for microbe metabolism, invasion-front toads have higher abundance of bacterial functions in factor 420 biosynthesis, critical to bacterial metabolism and mediating important redox transformations involved in bacterial persistence, antibiotic biosynthesis, pro-drug activation, and methanogenesis [60].

We also detected geographic variation in bacterial functional pathways that contribute to host health. Invasion-front toads exhibited bacterial functions beneficial to host health and immunity: (i) phosphopantothenate biosynthesis (involved in bacterial production of coenzyme A [61]); and (ii) superpathway of sialic acids biosynthesis (involved in immunity including acting as host receptors and pathogen decoys for viruses and bacteria [62] and especially critical for preventing neural tissue damage [63]). Despite this abundance of health-promoting bacterial functions, these toads may also face health challenges. Invasion-front toad bacteria had a higher abundance of urate biosynthesis function (urate biosynthesis/inosine 5’-phosphate degradation, the only significantly different one out of the top 30 abundant functions), which affects serum urate levels [64]. High levels of urate can result in the formation of needle-like crystals of urate in the joints (gout), perhaps related to severe spinal arthritis in invasion-front cane toads [65].

### Associations between lungworms and host gut bacteria

Pathogens and parasites impact the composition of the host microbiota and can modify host behaviour in a manner that improves parasite transmission and survival [66–68]. Lungworms can affect cane toad locomotor performance and reduce host endurance, curtailing oxygen supply from infected lungs [69]. Lungworms also can alter a cane toad’s thermal preference and manipulate the timing and location of defecation, thereby enhancing lungworm egg production and larval survival [22]. Lungworms are reported to lag behind their host on the invasion-front by 2-3 years [27] and to affect righting behaviour (prolongs righting time [70]). In the current study, although we collected invasion-front toads in recently invaded areas, we found no difference in lungworm presence or intensity between the invasion-front *versus* range-core toads, nor did we find behavioural differences in lungworm-infected *versus* uninfected toads.

Infection by parasitic lungworms was associated with differences in gut bacteria. Here, the direction of causation is less ambiguous than is the case for geographic variation in the gut bacteria. It seems unlikely that a toad’s bacteria affect its probability of carrying adult lungworms, although bacterial-driven differences in habitat selection might create such a link. Instead, we suggest that the presence of lungworms induces a shift in gut bacteria. Consistent with that hypothesis, experimental trials have shown that lungworms modify gastric function in their hosts, changing the volume and consistency of faeces produced in ways that enhance survival of larval lungworms [22]. Shifts in the microbiome inside the gut might be either causes or consequences of that shift in gastric function. Moreover, *C. elegans* are known to prefer specific bacterial foods [71], suggesting that lungworm larvae may also feed selectively on bacteria in the gut, generating differences in bacterial communities between lungworm-infected toads *versus* non-infected conspecifics. Additionally, gut bacteria may affect lungworms via microbiome-induced shifts in host immunity [72].

### Associations between host behaviours and gut bacteria

Interestingly, behaviours including righting effort likelihood and righting time were associated more closely with predicted gut bacterial functions than with bacterial taxonomic composition. Multiple identified taxa may share the same bacterial function, or one taxon may contribute to multiple bacterial functions, obscuring the relationship between host behaviour and bacterial taxonomic composition. Nonetheless, these relationships we found between gut bacterial function and righting behaviours may be related to toad health and/or rearing conditions. A dampened stress response (lower corticosterone levels) in invasion-front toads [73] could result from higher abundance of bacterial functions beneficial to host health and immunity, especially the superpathway of sialic acids biosynthesis [63]. Further, invasion-front toads are more reluctant to flee in simulated predation trials [74]. Dampened stress responses can be related to more exploratory behaviour [56], and to greater dispersal ability [26]. Rearing conditions also affect righting behaviour [17]. Although manipulative studies are needed to clarify causal relationships between stress responses, proactive behaviours, and gut bacterial functions, our results indicate that host behaviour and gut bacterial functions are related, suggesting that gut bacteria may be an important driver of invasion.

Our study has identified patterns rather than testing alternative hypotheses about underlying causal processes. To clarify causal mechanisms underlying the geographic divergence in gut bacteria across the toads’ Australian range, future studies could use reciprocal transplantation to examine if (and how) their gut bacteria respond to novel environmental conditions. Breeding these animals, and raising their offspring under common-garden conditions, could reveal the degree to which a toad’s gut bacteria is driven by host genetics *versus* their rearing conditions [75,76]. To clarify the hypothesis that changes in gut bacteria mediate the ability of lungworm parasites to modify host gut function, we could implant colon contents from infected into uninfected toads. In short, our discovery of strong associations between gut bacteria and important facets of toad ecology provides the opportunity to move to hypothesis-testing experimental studies.

Our research illustrates that during invasion, as a species expands across a novel and variable landscape, a complex relationship between host behaviour, its parasite community, and its microbiome may unfold. A clearer understanding of these relationships and how they influence the rate of expansion are key to understanding the role of the holobiont during invasion [77]. Such studies also will advance our understanding of co-evolution and may facilitate innovative approaches to invasive species management.

## Ethics

Approved by University of Adelaide Animal Ethics Committee (S-2018-056).

## Author Contributions

Designed research: JZ, TMN, CRL, SJZ, LAR; performed research: JZ, CRL, GWF, KS, LAR; analyzed data: JZ, TMN, CRL, SJZ, KS, LAR; drafted manuscript: JZ, RS, TMN, CRL, SJZ, GWF, KS, LAR.

## Acknowledgments

We thank Dr. Alice Russo and Ms. Rita Kurpiewski for field assistance, and Dr. Eve Slavich (UNSW Stats Central) for statistical advice.

## Data Availability Statement

Supplementary methods and results available online. Code available at: https://github.com/jiazhou0116/gut-microbiome-analyses-2. Sequence data are available in NCBI Sequence Read Archive (PRJNA670039). Raw ecological data available on Dryad (doi:10.5061/dryad.v15dv41tw).

