## Supplementary Materials for "The gut bacteria of an invasive amphibian respond to the dual challenges of range-expansion and parasite attack"

### Supplementary Methods

#### *Sample collection*

Justification of using colon contents

We selected the colon for analysis of bacteria because host behaviour appears to be more closely linked to the bacteria of the colon rather than that of the small intestine. The microbial community in the colon can ferment complex carbohydrates, such as dietary fibre, into short chain fatty acids (SCFAs) with neuroactive properties [1]. Microbial species in the colon can change brain metabolites (in pigs, *Ruminococcus* spp. can influence brain N-acetylaspartate through the mediation of serum cortisol [2]. In cane toads, reactive nerve cell bodies are more common in the submucosa of colon than in the small intestine [3]. Also, colon bacteria may have more effect on hosts than bacteria in other parts of the digestive system. Following regular nutrient provisioning, bacterial communities replicate quickly and reach the stationary phase (the point at which the maintenance of the bacterial population size reaches equilibrium) after only 20 minutes in the colon *versus* more than 2.2 hours in other parts of the digestive system. bacteria also have longer transit times in the colon (10-60 hours *versus* less than 3 hours in all other digestive organs [4]), increasing their opportunity to impact the host.

#### *Analyses of morphological and ecological data*

Firstly, we conducted tests to exclude highly correlated variables. We used t-tests to compare SUL between regions and between infected and non-infected toads with a consideration that standard deviations between invasion-front and range-core individuals are not equal.

We compared count and occurrence data (i.e., absence=0, presence=1) for parasite infection and behavioural traits between regions. To analyse regional differences in parasite occurrence data, we used a GLM with logistic regression (R version 4.0.2 [5]) and with a negative binomial regression *glm.nb* in the “MASS” package [6] to analyse over-dispersed count data, when the conditional variance exceeded the conditional mean.

We compared count data for struggle scores, righting effort and righting time in infected *versus* non-infected toads. To analyse struggle likelihood and righting effort likelihood (i.e., absence=0, presence=1) with respect to infection status, we used a GLM with logistic regression and with a negative binomial regression *glm.nb* in the "MASS" package to analyse over-dispersed count data, when the conditional variance exceeded the conditional mean.

***Molecular methodology***

We extracted microbial DNA from colon contents following the manufacturer's protocol for the DNeasy PowerSoil kit (Qiagen) and prepared 16S rRNA amplicon libraries using guidelines for the Illumina MiSeq System. We used Zymo isolated DNA (D6305) as a community positive control and MilliQ water as a PCR negative control to identify environmental contamination during the library preparation process [7]. Libraries were prepared based on the hypervariable (V3-V4) region of the 16S rRNA gene from each sample using primers 341F (5' – **TCGTCGGCAGCGTCAGATGTGTATAAGAGACAG**CCTACGGGNGGCWGCAG-3') and 785R (5'-**GTCTCGTGGGCTCGGAGATGTGTATAAGAGACAG**GACTACHVGGGTATCTAATCC-3') [8]. The library preparation details have been reported previously [9]. The libraries were sequenced at the Ramaciotti Centre for Genomics (University of New South Wales, Kensington, Sydney) on the Illumina MiSeq platform targeting 2x300bp paired-end sequence reads.

We downloaded demultiplexed FASTQ data from Illumina's BaseSpace cloud storage, then created an amplicon sequence variants (ASVs) abundance table using the open-source QIIME2 pipeline [10]. Demultiplexed sequence counts from samples and the positive control ranged between 138,989 and 305,662; the count of the PCR negative control was 67,506. The DADA2 pipeline [11], implemented in QIIME2, was used to filter and trim the first 20 bases from each read and truncate sequences to 200 bases. The remaining sequences were dereplicated, then forward/reverse reads were merged, chimeras were removed, and finally ASVs were generated for downstream analysis [11]. After quality filtering, reads from colon samples and the positive control ranged between 103,245 and 245,059 counts; the PCR negative control yielded 6,727 reads. The taxonomic assignment of ASVs was performed using Greengenes version 13_8 [12]. Data were pruned to remove representatives classified to Archaea (N = 28), chloroplast (N = 17), mitochondria (N = 186), and 151 unassigned (“kingdom”) ASVs implemented in the package ‘phyloseq’ [13]. We also removed the ASVs with prevalence of less than four, which makes the logged counts per sample more evenly distributed. The remaining 9,878 taxa were classified to the Kingdom Bacteria with 62.62% assigned to phylum level and 39.65% assigned to family level.

***Molecular data analyses***

To estimate alpha diversity (within individuals), we calculated observed ASVs [12], evenness [14], and Shannon [15] indices using QIIME2. Because the Shannon estimates of alpha diversity were highly correlated with those from the other indices (Spearman correlation > 0.8), we used the former for downstream analyses. We used a GLM model to compare Shannon indices across localities, including SUL as a covariate.

In order to explore differences in bacterial taxa across sampling sites (beta diversity), we first calculated the Core50 gut community [16] as follows: the ASV table was filtered to include only the ASVs present in a minimum of 50% of individual toads from each site. This calculation was performed separately for three sites from invasion-front toads: Kununurra (gut Core50: n = 111 ASVs), Old Theda (gut Core50: n = 118 ASVs), and Mary Pool (gut Core50: n = 129 ASVs); three sites from range-core toads: Rossville (gut Core50: n = 148 ASVs), Croydon (gut Core50: n = 86 ASVs), and Lucinda (gut Core50: n = 117 ASVs). Then we subsequently compiled filtered ASVs of six sites to avoid excluding ASVs that may be specific to only one site. In combination, the gut Core50 contained 325 unique ASVs. We visualized relative abundance of bacteria (abundance >2%) in different sample types, classified to the phylum and family level.

We then transformed the Core50 data set using a Hellinger transformation implemented in the package “microbiome” [17] in R and generated a Bray-Curtis dissimilarity matrix. We visualized this using Principal Coordinate Analysis (PCoA) plots using commands from the package “Phyloseq” [13]. We used the *adonis* command from the package “Vegan” to perform permutational multivariate analysis of variance (perMANOVA) to check whether the bacterial communities of toads from each region were significantly different. We used the command *betadisper* in the package “Vegan” [18] to check the homogeneity of group variances, an assumption of perMANOVA. After finding significant differences between invasion-front and range-core toads, we performed pairwise comparisons between all sampling sites using the command *pairwise.perm.manova* function in “RVAideMemoire” package with the Wilks Lambda [19]. We corrected for multiple testing [20] using the Hochberg procedure [21].

We used PICRUST2 [22] to predict bacterial functions for Core50 ASVs and generated pathway abundances considering the taxonomic contributions of ASVs. We removed all pathways with less than 0 prevalence and 474 out of 484 pathways remained for analysis. Then we used a Hellinger transformation and calculated and plotted Bray-Curtis dissimilarity matrix through package “Phyloseq”. We performed permutational multivariate analysis of variance (perMANOVA) to check whether the pathway abundance of toads from each location and site were significantly different and we used the command *betadisper* to check the homogeneity of group variances.

To identify differences in ASVs and predicted bacterial functions between range-core and invasion-front toads, we performed differential abundance testing using the function *DESeq* in the package “DESeq2” [23]. We report significant differences as the log fold change of a taxa in range-core toads *versus* those from the invasion-front. We set ASVs and bacterial functions abundance in invasion-front toads as the control group.

To identify associations between host characteristics (including infection with parasites) with bacterial communities and predicted bacterial functions, statistical analysis was conducted using the function envfit in the package “Vegan” [18] on Bray-Curtis dissimilarities. Bray-Curtis dissimilarities were visualized with non-metric multidimensional scaling (nMDS) plots. Data on host characteristics were fitted to the ordination plots using the function envfit.

After analyzing all variables individually, we also conducted distance-based redundancy analysis (dbRDA) combining all host characteristics to identify relationships between variables with respect to their impact on gut bacterial community. dbRDA performs constrained ordination directly on a distance or dissimilarity matrix with the function *capscale* in the “Vegan” package in R. Bray-Curtis dissimilarity matrixes of ASVs and predicted bacterial functions were ordinated and the results were analyzed using redundancy analysis with constraining variables that were not highly correlated (Spearman correlation coefficient < 0.85 and > -0.85) to estimate their explanatory proportion.

We explicitly tested whether lungworms affected bacterial communities. We used the *adonis* command from the package “Vegan” to perform permutational multivariate analysis of variance (perMANOVA) using Bray-Curtis distance metrics to check whether the bacterial communities of toads from infected *versus* non-infected toads were significantly different. We set bacterial Bray-Curtis distances as response, and region, count and occurrence of lungworm as factors. We used the command *betadisper* in the package "Vegan" to check the homogeneity of group variances, an assumption of perMANOVA.

### Supplementary Results

#### *Quality of DNA sequence data*

The number of observed species in the PCR negative control (N = 36) was lower than in toad colon samples (mean = 336, SD = 137.8), indicating minimal contamination throughout the library preparation processes. The number of observed species (N = 19) in the community positive control was also lower than in the experimental samples. The eight species used as bacterial community standards constituted more than 99.8% of ASVs from our sequenced community positive control, indicating minimal PCR bias in our library preparation process.

#### *Comparison of bacterial composition and diversity within individuals*

The alpha diversity (diversity within individual toads) did not differ significantly between range-core *versus* invasion-front toads: observed species (*wilcox.test*, p-value = 0.631), Pielou's evenness (*wilcox.test*, p-value = 0.307), Shannon (*wilcox.test*, p-value = 0.230). The Lucinda site had the highest Shannon diversity, followed by Old Theda, Mary Pool, Rossville, Croydon and Kununurra. Shannon diversity in individual samples was not significantly correlated with measured behavioural traits (*glm*, covariate = SUL, p-values > 0.050).

#### *Relative abundance of core50 bacteria in the colon*

A total of 89.23% ASVs were assigned to the level of phylum and dominant phyla (average abundance > 2%) were Firmicutes, Bacteroidetes, Proteobacteria, Fusobacteria, and Verrucomicrobia (Figure S1). We discuss the bacterial functional significance of these groups in the discussion. Across all individuals, we found that: (1) the dominant phyla were Firmicutes (average abundance = 47.89%, SD = 19.25%) and Bacteroidetes (average abundance = 39.05%, SD = 19.26%); (2) the phylum Bacteroidetes was present in all except two individual toads (Toad 14, 21) from the invasion-front and two toads (Toad 43, 48) from the range-core; and (3) one invasion-front toad (Toad 21) showed a much higher abundance of Proteobacteria (77.35%) than did other individuals (average abundance = 5.13%, SD = 7.99%).

A total of 70.77% ASVs were assigned to the family level and dominant families (average abundance > 2%) included Bacteroidaceae, Lachnospiraceae, Porphyromonadaceae, Clostridiaceae, Enterobacteriaceae, Erysipelotrichaceae, Fusobacteriaceae, Bacillaceae, Ruminococcaceae, Verrucomicrobiaceae, Turicibacteraceae, and Veillonellaceae. Toad 21, with a high abundance of Proteobacteria, also had more Enterobacteriaceae (77.34%) than did any other individuals (average abundance = 4.55%, SD = 7.86%).

A total of 40.62% ASVs were assigned to the genus level and dominant genera (average abundance > 2%) included *Bacteroides*, *Parabacteroides*, *Clostridium*, *Epulopiscium*, *Akkermansia*, *Turicibacter*, *Bacillus*.

***Relative abundance of core50 predicted bacterial functions in the colon***

The six most abundant pathways were pentose phosphate pathway (non-oxidative branch), adenosine deoxyribonucleotides de novo biosynthesis II, guanosine deoxyribonucleotides de novo biosynthesis II, pyruvate fermentation to isobutanol (engineered), Calvin-Benson-Bassham cycle, and adenine and adenosine salvage III (Figure 3).

### Supplementary Tables

**Table S1.** Sample collection sites and dates.

| Regions | Sites | Coordinates | Sample size | Collection dates |
|---|---|---|---|---|
| Western Australia 'invasion-front' | Kununurra | 15.776566° S, 128.744293° E | 10 | 20/11/2018 |
| | Old Theda | 14.790795° S, 126.497624° E | 10 | 21/11/2018 |
| | Mary Pool | 18.72528° S, 126.870096° E | 10 | 25/11/2018 |
| Queensland (QLD) 'range-core' | Rossville | 15.697069° S, 145.254385° E | 10 | 4/12/2018 |
| | Croydon | 18.207536° S, 142.245702° E | 10 | 5/12/2018 |
| | Lucinda | 18.530149° S, 146.331264° E | 10 | 6/12/2018 |

**Table S2.** Comparison of host characteristics and behavioural traits between range-core and invasion-front toads.

| Variables | t. test | GLM p-values (^SUL as covariate) | |
|---|---|---|---|
| | SDs not equal | glm.nb | binomial |
| *Host characteristics* | | | |
| Body length (SUL) | 0.014* | | |
| Lungworms^ | | 0.860 | |
| Occurrence of lungworms^ | | | 0.077 |
| Gut parasites^ | | 0.818 | |
| Occurrence of gut parasites^ | | | 0.604 |
| *Behavioural traits* | | | |
| Struggle score^ | | 0.002** | |
| Struggle likelihood^ | | | 0.008** |
| Righting effort^ | | 0.320 | |
| Righting effort likelihood^ | | | 0.036* |
| Righting time^ | | 0.507 | |

Note: Negative binomial regression (glm.nb) was used for over-dispersed count data, that is when the conditional variance exceeds the conditional mean. Significance codes: 0 '***' ≤0.001 '**' ≤0.01 '*' ≤0.05. SUL: snout urostyle length (body length). SDs not equal: true difference in means of invasion-front and range-core is not equal to 0.

**Table S3.** Mean and SD of host characteristics and behavioural traits of cane toads in this study.

| Variables | Range-core | | Invasion-front | | Lungworm-infected | | Lungworm-free | | Australia | |
|---|---|---|---|---|---|---|---|---|---|---|
| | mean | SD | mean | SD | mean | SD | mean | SD | mean | SD |
| *Host characteristics* | | | | | | | | | | |
| SUL | 96.24 | 12.50 | 103.51 | 9.42 | 101.12 | 12.59 | 98.55 | 10.43 | 99.88 | 11.57 |
| Body weight | 98.90 | 38.19 | 143.87 | 49.63 | 122.42 | 53.58 | 120.28 | 45.46 | 121.38 | 49.41 |
| Lungworms | 2.33 | 3.01 | 2.97 | 7.02 | - | - | - | - | 2.65 | 5.37 |
| Occurrence of lungworms | 0.60 | 0.50 | 0.43 | 0.50 | - | - | - | - | 0.52 | 0.50 |
| Gut parasites | 1.55 | 3.53 | 4.40 | 6.07 | 2.34 | 4.60 | 4.52 | 6.14 | 3.26 | 5.35 |
| Occurrence of gut parasites | 0.35 | 0.49 | 0.60 | 0.50 | 0.41 | 0.50 | 0.62 | 0.50 | 0.50 | 0.51 |
| *Behavioural traits* | | | | | | | | | | |
| Struggle scores | 2.63 | 2.53 | 2.10 | 4.54 | 1.84 | 2.50 | 2.93 | 4.56 | 2.37 | 3.65 |
| Struggle likelihood | 0.84 | 0.37 | 0.28 | 0.43 | 0.55 | 0.51 | 0.59 | 0.50 | 0.57 | 0.50 |
| Righting effort | 2.20 | 2.20 | 2.20 | 4.57 | 2.06 | 2.17 | 2.34 | 4.65 | 2.20 | 3.56 |
| Righting effort likelihood | 0.73 | 0.45 | 0.50 | 0.51 | 0.68 | 0.48 | 0.55 | 0.51 | 0.62 | 0.49 |
| Righting time (sec) | 24.83 | 47.36 | 12.93 | 34.03 | 20.35 | 43.32 | 17.31 | 39.77 | 18.88 | 41.32 |

**Table S4.** Comparison of host characteristics and behavioural traits between lungworm-infected and lungworm-free toads.

| Variables | t. test | GLM p-values (^SUL as covariate) | |
|---|---|---|---|
| | SDs not equal | glm.nb | binomial |
| *Host characteristics* | | | |
| Body length (SUL) | 0.393 | | |
| *Behavioural traits* | | | |
| Struggle score^ | | 0.229 | |
| Struggle likelihood^ | | | 0.906 |
| Righting effort^ | | 0.737 | |
| Righting effort likelihood^ | | | 0.316 |
| Righting time^ | | 0.771 | |

Note: Negative binomial regression (glm.nb) was used for over-dispersed count data, that is when the conditional variance exceeds the conditional mean. SUL: snout urostyle length (body length). SDs not equal: true difference in means of invasion-front and range-core is not equal to 0.

**Table S5.** Pairwise comparisons of cane toad gut bacterial composition beta diversity using permutation MANOVAs on a Bray Curtis distance matrix of Core50 bacteria.

| | Kununurra | Old Theda | Mary Pool | Rossville | Croydon |
|---|---|---|---|---|---|
| Old Theda | 2e-04 | - | - | - | - |
| Mary Pool | 2e-04 | 2e-04 | - | - | - |
| Rossville | 2e-04 | 2e-04 | 2e-04 | - | - |
| Croydon | 2e-04 | 2e-04 | 2e-04 | 2e-04 | - |
| Lucinda | 2e-04 | 2e-04 | 2e-04 | 2e-04 | 2e-04 |

P value adjustment method: Hochberg (1988)

**Table S6** Differences in abundance of specific bacterial ASVs in large intestine from range-core cane toads versus invasion-front cane toads (only including those that classified to Family level).

| ASV ID | baseMean | log2 Fold-Change | lfcSE | stat | pvalue | padj | Phylum | Class | Order | Family | Genus | Species |
|---|---|---|---|---|---|---|---|---|---|---|---|---|
| 6c58d45d4b8ff8cdecd8e9d1fb4c5a69 | 1487.697 | -3.250 | 1.037 | -3.134 | 1.727E-03 | 3.712E-03 | Proteo-bacteria | Gammaproteo-bacteria | Entero-bacteriales | Entero-bacteriaceae | NA | NA |
| 7557c00fea8f70259b8904b3b3d76880 | 895.763 | 2.531 | 0.993 | 2.550 | 1.078E-02 | 2.176E-02 | Firmicutes | Clostridia | Clostridiales | Lachno-spiraceae | Epulo-piscium | NA |
| cbdb93196f42014d5b0ab3ab42c17f88 | 527.943 | 3.495 | 1.155 | 3.026 | 2.475E-03 | 5.270E-03 | Proteo-bacteria | Gammaproteo-bacteria | Entero-bacteriales | Entero-bacteriaceae | NA | NA |
| 6419daf56085a3de7da0ea88506dc9fd | 764.777 | 1.842 | 0.642 | 2.870 | 4.105E-03 | 8.583E-03 | Firmicutes | Clostridia | Clostridiales | Lachno-spiraceae | NA | NA |
| a267538570c1388d346a727261fe91be | 172.240 | -24.844 | 1.694 | -14.662 | 1.126E-48 | 1.439E-47 | Bacter-oidetes | Bacteroidia | Bacteroidales | Bacteroid-aceae | Bacter-oides | NA |
| 0162cd21d987db84a48ea32e8cce0eb7 | 56.217 | -25.854 | 2.426 | -10.657 | 1.621E-26 | 1.130E-25 | Bacter-oidetes | Bacteroidia | Bacteroidales | Bacteroid-aceae | Bacter-oides | NA |
| e46a7bcff7461ee34f3b6b0d69df6d19 | 631.347 | -2.136 | 0.956 | -2.235 | 2.544E-02 | 4.757E-02 | Bacter-oidetes | Bacteroidia | Bacteroidales | Bacteroid-aceae | Bacter-oides | NA |
| e5cb7a6bb59f52f361cf3760b9f9ff4f | 104.997 | -26.595 | 2.275 | -11.689 | 1.454E-31 | 1.393E-30 | Bacter-oidetes | Bacteroidia | Bacteroidales | Bacteroid-aceae | Bacter-oides | NA |
| 80222d96e87adadb78c4b491e56a576b | 421.000 | -2.742 | 1.232 | -2.225 | 2.605E-02 | 4.832E-02 | Firmicutes | Clostridia | Clostridiales | Veillonell-aceae | NA | NA |
| 492e18555f4701f64bc74822c9b95b9a | 105.234 | -26.732 | 2.671 | -10.008 | 1.406E-23 | 6.879E-23 | Bacter-oidetes | Bacteroidia | Bacteroidales | Porphyro-monadaceae | NA | NA |
| 86290c20aa914c92a60c3b1928ef8552 | 84.244 | -26.423 | 2.656 | -9.950 | 2.511E-23 | 1.133E-22 | Firmicutes | Clostridia | Clostridiales | Lachno-spiraceae | NA | NA |
| 8c7ded70bf5271c289232d20c8565f34 | 151.400 | -27.231 | 1.391 | -19.583 | 2.140E-85 | 1.231E-83 | Bacter-oidetes | Bacteroidia | Bacteroidales | Bacter-oidaceae | Bacter-oides | ovatus |
| a4a53a3ed7fc3c92875b6d43d0f59a1b | 233.009 | -27.716 | 2.735 | -10.133 | 3.928E-24 | 2.101E-23 | Firmicutes | Clostridia | Clostridiales | Lachno-spiraceae | NA | NA |
| 780e6a8bc3c4294880771d06eb0d14cf | 25.306 | -24.247 | 2.595 | -9.343 | 9.389E-21 | 2.879E-20 | Firmicutes | Clostridia | Clostridiales | Lachno-spiraceae | NA | NA |
| 6902008369ce59136e20a96f2acd5e8b | 60.760 | -25.130 | 1.437 | -17.491 | 1.691E-68 | 4.322E-67 | Firmicutes | Erysipelotrichi | Erysipelo-trichales | Erysipelo-trichaceae | cc_115 | NA |
| a34ef14f8bbed084541dbbc7738bbcf6 | 81.138 | -26.363 | 1.535 | -17.179 | 3.800E-66 | 7.283E-65 | Firmicutes | Clostridia | Clostridiales | Lachno-spiraceae | NA | NA |
| 7b8c080caae57d2f824112c9a7fc3a6a | 53.462 | -25.778 | 2.069 | -12.460 | 1.230E-35 | 1.414E-34 | Bacter-oidetes | Bacteroidia | Bacteroidales | Bacteroid-aceae | Bacter-oides | NA |
| ef7cc9c06b4d9a3393bcb9e79db07c61 | 20.440 | -24.444 | 2.583 | -9.464 | 2.972E-21 | 9.493E-21 | Firmicutes | Clostridia | Clostridiales | Lachno-spiraceae | NA | NA |
| 54e630043d77fd8382fa7f3b6ed035fa | 13.532 | -22.453 | 2.910 | -7.716 | 1.199E-14 | 2.759E-14 | Bacter-oidetes | Bacteroidia | Bacteroidales | [Odori-bacteraceae] | Odori-bacter | NA |
| b5b8abbd375f4b9fcdc0af60d2777c99 | 30.332 | -24.987 | 2.378 | -10.509 | 7.891E-26 | 4.776E-25 | Bacter-oidetes | Bacteroidia | Bacteroidales | Bacteroid-aceae | Bacter-oides | NA |
| e1daff26a7f4ebcdd446769acc793647 | 26.210 | -24.542 | 2.205 | -11.128 | 9.163E-29 | 7.526E-28 | Firmicutes | Erysipelotrichi | Erysipelo-trichales | Erysipelo-trichaceae | [Eubac-terium] | dolichum |
| b10a19c6f57b7ef748a93f0a36046246 | 53.135 | -25.591 | 1.476 | -17.341 | 2.304E-67 | 4.818E-66 | Bacter-oidetes | Bacteroidia | Bacteroidales | Bacteroid-aceae | Bacter-oides | uniformis |

| | | | | | | | | | | | | |
|---|---|---|---|---|---|---|---|---|---|---|---|---|
| 3af5a570a8cbca4da5182555f5b8a257 | 6.010 | -6.051 | 2.631 | -2.300 | 2.143E-02 | 4.142E-02 | Firmicutes | Clostridia | Clostridiales | Veillonell-aceae | NA | NA |
| 3fbd34428a142daa4c75dc669aa7d8ce | 69.508 | -25.872 | 2.643 | -9.787 | 1.280E-22 | 5.257E-22 | Firmicutes | Clostridia | Clostridiales | Lachno-spiraceae | NA | NA |
| 60a58022cb997004475c70088245e826 | 4.819 | -5.733 | 2.390 | -2.399 | 1.643E-02 | 3.203E-02 | Actino-bacteria | Coriobacteriia | Corio-bacteriales | Corio-bacteriaceae | NA | NA |
| ed7f5ceb297ad51ce5945e0f55a98027 | 14.344 | -23.948 | 2.860 | -8.373 | 5.610E-17 | 1.330E-16 | Proteo-bacteria | Gammaproteo-bacteria | Entero-bacteriales | Entero-bacteriaceae | NA | NA |
| 4d05437a5f453059f791e488873c77e4 | 69.534 | -26.157 | 2.640 | -9.907 | 3.892E-23 | 1.689E-22 | Firmicutes | Clostridia | Clostridiales | Lachno-spiraceae | NA | NA |
| 39c8a7e2acbaab44f409f03f267e754b | 546.170 | 5.515 | 1.027 | 5.371 | 7.819E-08 | 1.746E-07 | Firmicutes | Bacilli | Bacillales | Bacillaceae | Bacillus | NA |
| 44f67bdfbd628b322769918ba839915a | 248.837 | -27.923 | 1.423 | -19.620 | 1.041E-85 | 7.981E-84 | Bacter-oidetes | Bacteroidia | Bacteroidales | Porphyro-monadaceae | Parabac-teroides | NA |
| 8ee379f05f6903bf6d292d30a9fd4395 | 96.668 | -26.609 | 2.254 | -11.807 | 3.597E-32 | 3.597E-31 | Bacter-oidetes | Bacteroidia | Bacteroidales | Bacteroid-aceae | Bacteroides | NA |
| 174d67bd393815c98bb437be24448376 | 277.450 | -28.094 | 1.527 | -18.404 | 1.223E-75 | 4.018E-74 | Bacter-oidetes | Bacteroidia | Bacteroidales | Porphyro-monadaceae | Parabac-teroides | distasonis |
| afb88164af28994054d9be391f1ad7af | 82.816 | -26.401 | 2.439 | -10.824 | 2.651E-27 | 2.032E-26 | Bacter-oidetes | Bacteroidia | Bacteroidales | Bacteroid-aceae | Bacter-oides | NA |
| 2b7a7d24b576f24d679fa5c353f0f7f3 | 26.501 | -24.806 | 2.589 | -9.580 | 9.714E-22 | 3.529E-21 | Verruco-microbia | Verruco-microbiae | Verruco-microbiales | Verruco-microbiaceae | Akker-mansia | NA |
| b7a2c1afa54353b8946b9d7c115c7a44 | 189.563 | 4.089 | 1.632 | 2.506 | 1.223E-02 | 2.424E-02 | Bacter-oidetes | Bacteroidia | Bacteroidales | Rikenellaceae | NA | NA |
| 8f53eb1f0dd7db73f3995c22fe4f2388 | 64.676 | -26.055 | 2.421 | -10.760 | 5.312E-27 | 3.818E-26 | Firmicutes | Erysipelotrichi | Erysipelo-trichales | Erysipelo-trichaceae | NA | NA |
| 5d7b261edd80dc99e0c4100e8db8cfdb | 198.905 | -27.622 | 1.505 | -18.351 | 3.217E-75 | 9.250E-74 | Bacter-oidetes | Bacteroidia | Bacteroidales | Bacteroid-aceae | Bacter-oides | ovatus |
| c8f8bb35d044c738a35039ae87bae185 | 11.424 | -23.621 | 2.586 | -9.136 | 6.509E-20 | 1.782E-19 | Bacter-oidetes | Bacteroidia | Bacteroidales | [Odori-bacteraceae] | Odori-bacter | NA |
| d081db749cdc490e77383b1059f85d8f | 50.480 | -25.709 | 2.422 | -10.613 | 2.599E-26 | 1.758E-25 | Firmicutes | Clostridia | Clostridiales | Lachno-spiraceae | Epulo-piscium | NA |
| b908275e7b7716ca1c353d1c7b844483 | 67.899 | -26.111 | 1.618 | -16.135 | 1.455E-58 | 2.390E-57 | Bacter-oidetes | Bacteroidia | Bacteroidales | Rikenellaceae | AF12 | NA |
| 2ee89c589341947cb387326928eb95e5 | 35.058 | -25.192 | 2.199 | -11.457 | 2.158E-30 | 1.985E-29 | Firmicutes | Erysipelotrichi | Erysipelo-trichales | Erysipelo-trichaceae | NA | NA |
| 590ca5ec1fa06013677ae3d8337043f3 | 36.909 | -24.294 | 2.870 | -8.466 | 2.541E-17 | 6.216E-17 | Bacte-roidetes | Bacteroidia | Bacteroidales | Rikenellaceae | NA | NA |
| e991093fcfddadae8044626080e8f7c6 | 51.056 | -25.469 | 1.697 | -15.008 | 6.516E-51 | 8.816E-50 | Firmicutes | Clostridia | Clostridiales | Clostridiaceae | Clostri-dium | NA |
| a1ba648cceb09bd2f407b369f3e64b55 | 38.587 | -24.619 | 2.611 | -9.428 | 4.180E-21 | 1.317E-20 | Bacter-oidetes | Bacteroidia | Bacteroidales | Bacteroid-aceae | Bacter-oides | NA |
| adc1e93b602d5cba53afff8f74915944 | 76.552 | -24.446 | 1.624 | -15.052 | 3.349E-51 | 4.814E-50 | Bacter-oidetes | Bacteroidia | Bacteroidales | Porphyro-monadaceae | Parabac-teroides | NA |
| f16dc2d38b6d9adf50e8870184112060 | 24.228 | -24.676 | 2.597 | -9.502 | 2.058E-21 | 6.861E-21 | Bacter-oidetes | Bacteroidia | Bacteroidales | Porphyro-monadaceae | Parabac-teroides | NA |
| e39d5eb9eb23d683073ca0dbbad1a672 | 38.087 | -25.313 | 2.393 | -10.576 | 3.835E-26 | 2.520E-25 | Bacter-oidetes | Bacteroidia | Bacteroidales | Bacteroid-aceae | Bacter-oides | NA |
| 04e5b39fb640a01af7d62efa921adf13 | 7.078 | -6.288 | 2.362 | -2.662 | 7.779E-03 | 1.598E-02 | Firmicutes | Clostridia | Clostridiales | Rumino-coccaceae | Oscillo-spira | NA |

| | | | | | | | | | | | | |
|---|---|---|---|---|---|---|---|---|---|---|---|---|
| 74f47914a81d27d19d24fda1ba90542a | 24.910 | -24.720 | 2.592 | -9.539 | 1.445E-21 | 4.960E-21 | Bacter-oidetes | Bacteroidia | Bacteroidales | Rikenellaceae | PW3 | NA |
| 045d65d76b036254cf0e59ec1717e221 | 18.821 | -23.120 | 2.588 | -8.934 | 4.093E-19 | 1.070E-18 | Bacter-oidetes | Bacteroidia | Bacteroidales | Rikenellaceae | NA | NA |
| 3b8ffd49ee49e9dd639b32985c8ec6fd | 8.374 | -23.192 | 2.603 | -8.909 | 5.138E-19 | 1.328E-18 | Firmicutes | Clostridia | Clostridiales | Rumino-coccaceae | NA | NA |
| 519b011405b6f41ac7727b8eb1f1b5b4 | 15.799 | -24.084 | 2.593 | -9.289 | 1.560E-20 | 4.660E-20 | Firmicutes | Clostridia | Clostridiales | Rumino-coccaceae | Anaero-filum | NA |
| 0936ab029dadd9cd26863bc1b3b0c2f2 | 11.643 | -23.650 | 2.372 | -9.971 | 2.042E-23 | 9.586E-23 | Bacter-oidetes | Bacteroidia | Bacteroidales | Rikenellaceae | NA | NA |
| ddf9de7466cd35a6d2af7858afec335c | 40.417 | -25.398 | 2.604 | -9.755 | 1.757E-22 | 6.966E-22 | Bacter-oidetes | Bacteroidia | Bacteroidales | Porphyro-monadaceae | NA | NA |
| 2a3d09d2028de2b116bd1cc7986e6ca6 | 5.716 | -5.979 | 2.637 | -2.267 | 2.337E-02 | 4.442E-02 | Bacter-oidetes | Bacteroidia | Bacteroidales | Bacteroid-aceae | NA | NA |
| c2972dcd4d3380ce07afc32b4d56732c | 25.773 | -24.297 | 2.596 | -9.358 | 8.161E-21 | 2.536E-20 | Bacter-oidetes | Bacteroidia | Bacteroidales | Bacteroid-aceae | Bacter-oides | NA |
| bcfce8cc1f1d41d4ca4291dc70ba36d4 | 7.713 | -23.077 | 2.610 | -8.841 | 9.482E-19 | 2.423E-18 | Firmicutes | Clostridia | Clostridiales | Rumino-coccaceae | Oscillo-spira | NA |
| a84952a043073200cd76472c25259e0a | 9.382 | -22.779 | 2.596 | -8.775 | 1.711E-18 | 4.279E-18 | Bacter-oidetes | Bacteroidia | Bacteroidales | [Odori-bacteraceae] | Odori-bacter | NA |
| 5dab6fb4a24683c16a17a1e865db6f3b | 14.446 | -23.955 | 2.362 | -10.142 | 3.612E-24 | 1.978E-23 | Firmicutes | Clostridia | Clostridiales | [Mogi-bacteriaceae] | NA | NA |
| 49d79c5f2707995019dbb234ac26f5ce | 13.774 | -23.893 | 2.586 | -9.241 | 2.440E-20 | 7.104E-20 | Firmicutes | Clostridia | Clostridiales | Pepto-coccaceae | NA | NA |
| 648b48ced2889e428f20630130d56c62 | 15.776 | -7.445 | 1.621 | -4.594 | 4.348E-06 | 9.617E-06 | Firmicutes | Clostridia | Clostridiales | Christen-senellaceae | NA | NA |
| 6c5022573f8f973316205803e3059526 | 11.567 | -23.649 | 2.592 | -9.123 | 7.338E-20 | 1.985E-19 | Bacter-oidetes | Bacteroidia | Bacteroidales | Rikenellaceae | NA | NA |
| a6f3f75daa5b4b143aefea057c3b0340 | 5.848 | -6.012 | 2.641 | -2.276 | 2.285E-02 | 4.379E-02 | Bacter-oidetes | Bacteroidia | Bacteroidales | Rikenellaceae | NA | NA |
| 539b882160c167bb0e9be429075a4ea9 | 13.083 | -23.821 | 2.585 | -9.216 | 3.088E-20 | 8.768E-20 | Firmicutes | Clostridia | Clostridiales | Rumino-coccaceae | NA | NA |
| adcee925b55ce9e5ace0c30909d39f37 | 7.374 | -6.346 | 2.622 | -2.420 | 1.552E-02 | 3.051E-02 | Firmicutes | Clostridia | Clostridiales | Rumino-coccaceae | NA | NA |
| 177eb3f471db964278cb7b238f78a184 | 836.171 | 6.493 | 1.187 | 5.470 | 4.488E-08 | 1.012E-07 | Firmicutes | Clostridia | Clostridiales | Lachno-spiraceae | Epulo-piscium | NA |
| f9bede316aab2d20a7f9fb07840e565d | 32.180 | -25.073 | 2.622 | -9.563 | 1.147E-21 | 4.059E-21 | Bacter-oidetes | Bacteroidia | Bacteroidales | Bacteroid-aceae | Bacter-oides | NA |
| 009da521e089ef304d1f66c8b04e28ea | 32.882 | -25.103 | 2.387 | -10.516 | 7.252E-26 | 4.508E-25 | Bacter-oidetes | Bacteroidia | Bacteroidales | Porphyro-monadaceae | Parabac-teroides | distasonis |
| fe086df68487400843a44fb4e7ab9f16 | 15.184 | -24.027 | 2.910 | -8.257 | 1.491E-16 | 3.500E-16 | Bacter-oidetes | Bacteroidia | Bacteroidales | Rikenellaceae | NA | NA |
| 8dd58bf584f5a402ff021b724147e395 | 17.603 | -24.238 | 2.027 | -11.954 | 6.158E-33 | 6.744E-32 | Bacter-oidetes | Bacteroidia | Bacteroidales | Porphyro-monadaceae | Parabac-teroides | NA |
| f2663b93e6b7afbc782d5134276f0fdd | 52.178 | -25.508 | 2.415 | -10.564 | 4.361E-26 | 2.786E-25 | Bacter-oidetes | Bacteroidia | Bacteroidales | Porphyro-monadaceae | Parabac-teroides | Distasonis |
| 1724290d764f85a42c9db7d7cb939846 | 60.986 | 5.546 | 1.240 | 4.471 | 7.786E-06 | 1.705E-05 | Proteo-bacteria | Deltaproteo-bacteria | Desulfo-vibrionales | Desulfovi-brionaceae | Bilophila | NA |
| 04411f2511ea9f914f865deb3927832e | 34.979 | -25.121 | 2.600 | -9.661 | 4.398E-22 | 1.686E-21 | Bacter-oidetes | Bacteroidia | Bacteroidales | Porphyro-monadaceae | Parabac-teroides | NA |

| | | | | | | | | | | | | |
|---|---|---|---|---|---|---|---|---|---|---|---|---|
| 47546847c4397857f887fff788f08f60 | 13.313 | -23.846 | 2.366 | -10.077 | 6.999E-24 | 3.577E-23 | Firmicutes | Clostridia | Clostridiales | Veillonell-aceae | NA | NA |
| 1a9aa68607bc39a575a4b7c81be479c0 | 88.182 | -26.481 | 2.097 | -12.629 | 1.464E-36 | 1.772E-35 | Firmicutes | Clostridia | Clostridiales | Veillonell-aceae | Phascol-arcto-bacterium | NA |
| a01dd3da1aafce98d380fee8b94331be | 49.548 | -25.679 | 2.631 | -9.762 | 1.640E-22 | 6.619E-22 | Bacter-oidetes | Bacteroidia | Bacteroidales | Porphyro-monadaceae | Parabac-teroides | NA |
| 605564c34134416f446b516ae52d0179 | 29.076 | -24.894 | 2.201 | -11.309 | 1.190E-29 | 1.052E-28 | Firmicutes | Erysipelotrichi | Erysipelo-trichales | Erysipelo-trichaceae | NA | NA |
| 16a88166845e1d751f122c0dcca98a5b | 9.769 | -23.406 | 2.595 | -9.020 | 1.887E-19 | 5.047E-19 | Bacter-oidetes | Bacteroidia | Bacteroidales | Bacteroid-aceae | Bacter-oides | Ovatus |
| 64991db060643dc3bdfb8c0fdf51e6e3 | 13.321 | -23.841 | 2.362 | -10.096 | 5.781E-24 | 3.022E-23 | Bacter-oidetes | Bacteroidia | Bacteroidales | Rikenellaceae | NA | NA |
| 62bc67a80f888b27125bc2690bf3ee51 | 49.278 | -24.095 | 2.626 | -9.176 | 4.474E-20 | 1.255E-19 | Bacter-oidetes | Bacteroidia | Bacteroidales | Bacteroid-aceae | Bacter-oides | Ovatus |
| da0b19878792e922e6b3f6ccc84f77c5 | 24.955 | -24.713 | 2.187 | -11.300 | 1.312E-29 | 1.118E-28 | Bacter-oidetes | Bacteroidia | Bacteroidales | Bacteroid-aceae | Bacter-oides | Fragilis |
| b940997b332c20ab631937981f61654b | 6.271 | -6.113 | 2.163 | -2.827 | 4.704E-03 | 9.747E-03 | Bacter-oidetes | Bacteroidia | Bacteroidales | Bacteroid-aceae | Bacter-oides | NA |
| a2397f0089a72cfa1395121f94fc49eb | 17.390 | -7.581 | 2.622 | -2.891 | 3.835E-03 | 8.091E-03 | Firmicutes | Bacilli | Bacillales | Bacillaceae | Bacillus | NA |
| 93d1b46d5dc3e1619eea42b1bff80f05 | 651.062 | 29.816 | 1.909 | 15.616 | 5.626E-55 | 8.626E-54 | Firmicutes | Bacilli | Bacillales | Bacillaceae | NA | NA |
| b9a467ef0617f4d815fa015fb0a2ffd4 | 88.833 | 26.980 | 2.909 | 9.273 | 1.802E-20 | 5.315E-20 | Bacter-oidetes | Bacteroidia | Bacteroidales | Bacteroid-aceae | Bacter-oides | Fragilis |
| 6904cbd6783c02626a2aba468a5640f3 | 275.743 | 28.641 | 1.283 | 22.319 | 2.403E-110 | 2.764E-108 | Bacter-oidetes | Bacteroidia | Bacteroidales | Bacteroid-aceae | Bacter-oides | NA |
| a51279dcacf45d515b7eaacc70b802a4 | 308.968 | 28.784 | 1.529 | 18.827 | 4.519E-79 | 2.079E-77 | Bacter-oidetes | Bacteroidia | Bacteroidales | Porphyro-monadaceae | Parabac-teroides | Distasonis |
| be967ee151c8743bda1042cadc340ed5 | 203.438 | 27.890 | 2.516 | 11.084 | 1.494E-28 | 1.185E-27 | Bacter-oidetes | Bacteroidia | Bacteroidales | Porphyro-monadaceae | Parabac-teroides | NA |
| 210d4d637761db63f3f16c30ee407979 | 49.483 | 26.289 | 2.649 | 9.923 | 3.319E-23 | 1.468E-22 | Bacter-oidetes | Bacteroidia | Bacteroidales | Bacteroid-aceae | NA | NA |
| 339e1c8994f7fbc0a23010fe97bce89d | 15.545 | 24.708 | 2.606 | 9.480 | 2.555E-21 | 8.394E-21 | Bacter-oidetes | Bacteroidia | Bacteroidales | Rikenellaceae | NA | NA |
| 75c0ea2b7d539613e42a006091e3f56b | 142.138 | 27.733 | 1.490 | 18.612 | 2.583E-77 | 9.903E-76 | Bacter-oidetes | Bacteroidia | Bacteroidales | Porphyro-monadaceae | Parabac-teroides | Distasonis |
| 1692e6cc71419edc280c6c756ba9e0af | 224.980 | 28.362 | 1.157 | 24.517 | 9.627E-133 | 2.214E-130 | Firmicutes | Bacilli | Bacillales | Bacillaceae | Bacillus | NA |
| fd304167835f6234bae704afa06f4614 | 47.280 | 26.218 | 2.624 | 9.991 | 1.673E-23 | 8.017E-23 | Firmicutes | Erysipelotrichi | Erysipelo-trichales | Erysipelo-trichaceae | Anaero-rhabdus | Furcosa |
| 95cc31da651c0d1d75c6a86ac1159f26 | 88.364 | 27.081 | 1.549 | 17.480 | 2.026E-68 | 4.659E-67 | Firmicutes | Bacilli | Bacillales | Bacillaceae | Bacillus | NA |
| 4aa5cfe66220aa950fbd6db3361a3a86 | 44.608 | 26.145 | 2.624 | 9.962 | 2.237E-23 | 1.029E-22 | Firmicutes | Erysipelotrichi | Erysipelo-trichales | Erysipelo-trichaceae | NA | NA |
| f7ded0075eb755c68df28bc168423431 | 46.193 | 8.931 | 1.400 | 6.379 | 1.779E-10 | 4.050E-10 | Bacter-oidetes | Bacteroidia | Bacteroidales | Bacteroid-aceae | Bacter-oides | NA |
| eb29bde72a89e554fd3871a4c957a6dd | 61.146 | 25.323 | 2.431 | 10.417 | 2.080E-25 | 1.196E-24 | Bacter-oidetes | Bacteroidia | Bacteroidales | Bacteroid-aceae | Bacter-oides | Ovatus |
| 9f6285cb058aa80b10a2a2ddae126f79 | 68.520 | 9.499 | 1.124 | 8.449 | 2.934E-17 | 7.104E-17 | Firmicutes | Clostridia | Clostridiales | Rumino-coccaceae | Oscillo-spira | NA |

| | | | | | | | | | | | | |
|---|---|---|---|---|---|---|---|---|---|---|---|---|
| 226d5e071abccfc78b7782c435775a73 | 36.171 | 24.333 | 2.392 | 10.172 | 2.636E-24 | 1.479E-23 | Bacter-oidetes | Bacteroidia | Bacteroidales | Bacteroid-aceae | Bacter-oides | NA |
| 7a2bb48b8b3e3e6a46d949d0992af1a6 | 19.531 | 25.015 | 2.386 | 10.486 | 1.006E-25 | 5.932E-25 | Firmicutes | Clostridia | Clostridiales | Rumino-coccaceae | Rumino-coccus | NA |
| fb42aec635a286734175e40ced11d984 | 8.315 | 23.853 | 2.611 | 9.137 | 6.420E-20 | 1.779E-19 | Firmicutes | Clostridia | Clostridiales | Lachno-spiraceae | NA | NA |
| 324cd8f38431f705cb73a98d824808dd | 5.821 | 5.942 | 2.634 | 2.256 | 2.407E-02 | 4.538E-02 | Firmicutes | Clostridia | Clostridiales | Rumino-coccaceae | Anaero-truncus | NA |
| 37d1958bb40faa4f43062c658209c5f9 | 6.393 | 6.077 | 2.391 | 2.542 | 1.102E-02 | 2.205E-02 | Firmicutes | Clostridia | Clostridiales | Rumino-coccaceae | Oscillo-spira | NA |
| 5f3dc837177b94869a0aff87c5714c43 | 22.672 | 25.218 | 2.865 | 8.801 | 1.351E-18 | 3.413E-18 | Proteo-bacteria | Gammaproteo-bacteria | Entero-bacteriales | Entero-bacteriaceae | NA | NA |
| fe8896769cf56f04c74c8d954ba40f46 | 28.131 | 25.496 | 2.602 | 9.797 | 1.154E-22 | 4.827E-22 | Proteo-bacteria | Gammaproteo-bacteria | Entero-bacteriales | Entero-bacteriaceae | NA | NA |
| 27b633e38e3296a88295f474cbe3b0a8 | 15.458 | 24.689 | 2.589 | 9.537 | 1.469E-21 | 4.968E-21 | Firmicutes | Clostridia | Clostridiales | Rumino-coccaceae | Oscillo-spira | NA |
| 2e507d213cc3c9d5a8f671095253f35c | 36.719 | 25.336 | 1.541 | 16.446 | 8.889E-61 | 1.573E-59 | Firmicutes | Clostridia | Clostridiales | Rumino-coccaceae | Oscillo-spira | NA |
| 0db2ae9cddc3b7aa17cbd7b31385af59 | 34.505 | 25.781 | 2.386 | 10.806 | 3.221E-27 | 2.390E-26 | Firmicutes | Clostridia | Clostridiales | Rumino-coccaceae | Oscillo-spira | NA |
| 314fa069eccc1d6240d8af01638d5b41 | 6.889 | 23.594 | 2.910 | 8.107 | 5.173E-16 | 1.202E-15 | Firmicutes | Clostridia | Clostridiales | Rumino-coccaceae | NA | NA |
| b0b496c9204914b969eafc30901252ed | 17.328 | 24.847 | 2.594 | 9.579 | 9.820E-22 | 3.529E-21 | Bacter-oidetes | Bacteroidia | Bacteroidales | Rikenellaceae | NA | NA |
| 0f179b8bfed9e2286ea5f352e27b157e | 17.755 | 24.889 | 2.592 | 9.603 | 7.796E-22 | 2.940E-21 | Bacter-oidetes | Bacteroidia | Bacteroidales | Bacteroid-aceae | Bacter-oides | fragilis |
| 3c064bab12d667f909b0272d9faefaad | 8.650 | 6.511 | 1.516 | 4.296 | 1.743E-05 | 3.782E-05 | Firmicutes | Clostridia | Clostridiales | Clostridiaceae | Clostri-dium | sub-terminale |
| bcd408786a42363c89b7950ad357a38c | 12.417 | 24.377 | 2.910 | 8.378 | 5.403E-17 | 1.295E-16 | Proteo-bacteria | Alphaproteo-bacteria | Rhodo-spirillales | Rhodo-spirillaceae | NA | NA |
| a8e25c5d1456422d37addd96abcbb11b | 100.317 | 27.251 | 2.291 | 11.897 | 1.228E-32 | 1.284E-31 | Bacter-oidetes | Bacteroidia | Bacteroidales | Bacteroid-aceae | Bacter-oides | NA |
| 927c28b2c30481130d754bd0c4f50b66 | 23.278 | 25.258 | 2.600 | 9.716 | 2.587E-22 | 1.009E-21 | Firmicutes | Clostridia | Clostridiales | Lachno-spiraceae | NA | NA |
| 4a15ae0b8939617699a4c1f30262992d | 43.607 | 25.037 | 2.887 | 8.672 | 4.228E-18 | 1.046E-17 | Bacter-oidetes | Bacteroidia | Bacteroidales | Bacteroid-aceae | Bacter-oides | NA |
| 005555dabe243422d4d02f4404ab0bcc | 35.081 | 25.761 | 2.619 | 9.835 | 7.925E-23 | 3.375E-22 | Bacter-oidetes | Bacteroidia | Bacteroidales | Bacteroid-aceae | Bacter-oides | NA |
| a71e588587585eeccb10b72cd277e3d1 | 10.543 | 24.164 | 2.595 | 9.312 | 1.250E-20 | 3.784E-20 | Firmicutes | Clostridia | Clostridiales | Rumino-coccaceae | Anaero-truncus | NA |
| 205652d10f1305824452dc4031a8a515 | 22.071 | 24.747 | 2.593 | 9.543 | 1.389E-21 | 4.842E-21 | Firmicutes | Clostridia | Clostridiales | Rumino-coccaceae | Oscillo-spira | NA |
| a8415a87a5673fb528660c8b990cfb40 | 7.006 | 23.604 | 2.632 | 8.968 | 3.025E-19 | 7.996E-19 | Firmicutes | Bacilli | Bacillales | Bacillaceae | NA | NA |
| b5c52517ccb11e5fab20a6e4e367efc2 | 16.701 | 24.801 | 2.588 | 9.583 | 9.408E-22 | 3.490E-21 | Bacter-oidetes | Bacteroidia | Bacteroidales | Rikenellaceae | NA | NA |
| 93bbe1e0eda851951753f40d62553f82 | 6.396 | 6.078 | 2.380 | 2.553 | 1.066E-02 | 2.171E-02 | Firmicutes | Clostridia | Clostridiales | Rumino-coccaceae | Oscillo-spira | NA |
| 2dbe579fb8ea4ed05be478d8d23cdeab | 12.384 | 24.040 | 2.602 | 9.240 | 2.474E-20 | 7.112E-20 | Firmicutes | Clostridia | Clostridiales | Rumino-coccaceae | Oscillo-spira | NA |

| | | | | | | | | | | | | |
|---|---|---|---|---|---|---|---|---|---|---|---|---|
| 91127a840605abfe8cf4a1bc1bc501b4 | 8.821 | 23.935 | 2.382 | 10.050 | 9.185E-24 | 4.592E-23 | Firmicutes | Clostridia | Clostridiales | Rumino-coccaceae | NA | NA |
| 3fa4268f5769ec9ca10bd42eec388a02 | 13.714 | 24.526 | 2.588 | 9.476 | 2.632E-21 | 8.526E-21 | Firmicutes | Clostridia | Clostridiales | Rumino-coccaceae | Oscillo-spira | NA |

**Table S7.** Pairwise comparisons using permutation MANOVAs on a Bray Curtis distance matrix of Core50 predicted bacterial functions.

| | Kununurra | Old Theda | Mary Pool | Rossville | Croydon |
|---|---|---|---|---|---|
| Old Theda | 0.298 | - | - | - | - |
| Mary Pool | 0.298 | 0.298 | - | - | - |
| Rossville | 0.009 | 0.298 | 0.298 | - | - |
| Croydon | 0.105 | 0.298 | 0.298 | 0.298 | - |
| Lucinda | 0.046 | 0.298 | 0.298 | 0.298 | 0.298 |

P value adjustment method: Hochberg (1988)

**Table S8.** Differences in abundance of specific predicted functions in large intestinal bacterial community from range-core cane toads versus invasion-front cane toads.

| Pathway names | predicted functional pathways | baseMean | log2Fold-Change | lfcSE | stat | pvalue | padj |
|---|---|---|---|---|---|---|---|
| AEROBACTINSYN-PWY | aerobactin biosynthesis | 263.716 | -2.211 | 0.543 | -4.070 | 4.702E-05 | 6.045E-04 |
| ANAEROFRUCAT-PWY | homolactic fermentation | 43046.792 | -0.110 | 0.035 | -3.180 | 1.474E-03 | 1.184E-02 |
| ARGDEG-PWY | superpathway of L-arginine, putrescine, and 4-aminobutanoate degradation | 896.212 | 1.533 | 0.567 | 2.705 | 6.840E-03 | 3.855E-02 |
| CENTFERM-PWY | pyruvate fermentation to butanoate | 10961.088 | 0.514 | 0.184 | 2.796 | 5.175E-03 | 3.119E-02 |
| CRNFORCAT-PWY | creatinine degradation I | 552.210 | 3.810 | 0.575 | 6.625 | 3.478E-11 | 1.677E-09 |
| FOLSYN-PWY | superpathway of tetrahydrofolate biosynthesis and salvage | 38619.204 | -0.166 | 0.039 | -4.272 | 1.937E-05 | 2.712E-04 |
| GALLATE-DEGRADATION -I-PWY | gallate degradation II | 241.732 | 2.376 | 0.536 | 4.434 | 9.238E-06 | 1.485E-04 |
| GALLATE-DEGRADATION-II-PWY | gallate degradation I | 242.098 | 2.363 | 0.527 | 4.487 | 7.219E-06 | 1.253E-04 |
| GLUCONEO-PWY | gluconeogenesis I | 46507.393 | -0.094 | 0.030 | -3.088 | 2.015E-03 | 1.482E-02 |
| GLYCOLYSIS | superpathway of glycolysis and Entner-Doudoroff | 46490.176 | -0.129 | 0.032 | -4.062 | 4.875E-05 | 6.045E-04 |
| HCAMHPDEG-PWY | 3-phenylpropanoate and 3-(3-hydroxy-phenyl) propanoate degradation to 2-oxopent-4-enoate | 228.844 | 2.278 | 0.560 | 4.066 | 4.782E-05 | 6.045E-04 |
| LACTOSECAT-PWY | lactose and galactose degradation I | 1003.482 | -0.896 | 0.331 | -2.710 | 6.726E-03 | 3.855E-02 |
| METHANOGENESIS-PWY | methanogenesis from H2 and CO2 | 5.656 | -3.540 | 0.725 | -4.882 | 1.048E-06 | 2.166E-05 |
| METHYLGALLATE-DEGRADATION-PWY | methylgallate degradation | 300.947 | 2.369 | 0.535 | 4.426 | 9.578E-06 | 1.485E-04 |
| NADSYN-PWY | NAD biosynthesis II (from tryptophan) | 503.798 | 3.689 | 0.481 | 7.666 | 1.777E-14 | 1.195E-12 |
| NAGLIPASYN-PWY | lipid IVA biosynthesis | 20574.919 | -0.510 | 0.191 | -2.677 | 7.428E-03 | 4.029E-02 |
| ORNARGDEG-PWY | superpathway of L-arginine and L-ornithine degradation | 896.212 | 1.533 | 0.567 | 2.705 | 6.840E-03 | 3.855E-02 |
| P108-PWY | pyruvate fermentation to propanoate I | 32200.249 | -0.303 | 0.113 | -2.680 | 7.364E-03 | 4.029E-02 |
| P164-PWY | purine nucleobases degradation I (anaerobic) | 29501.263 | 0.377 | 0.108 | 3.483 | 4.953E-04 | 4.576E-03 |
| P221-PWY | octane oxidation | 536.493 | -1.229 | 0.432 | -2.846 | 4.430E-03 | 2.787E-02 |
| P241-PWY | coenzyme B biosynthesis | 4.218 | -2.542 | 0.700 | -3.633 | 2.804E-04 | 2.830E-03 |
| P381-PWY | adenosylcobalamin biosynthesis II (late cobalt incorporation) | 18.306 | -2.832 | 0.634 | -4.470 | 7.840E-06 | 1.309E-04 |
| P42-PWY | incomplete reductive TCA cycle | 44742.461 | -0.269 | 0.085 | -3.149 | 1.637E-03 | 1.269E-02 |
| POLYISOPRENSYN-PWY | polyisoprenoid biosynthesis (E. coli) | 30996.681 | -0.282 | 0.077 | -3.666 | 2.466E-04 | 2.549E-03 |
| PWY-1541 | superpathway of taurine degradation | 173.948 | 1.788 | 0.579 | 3.088 | 2.012E-03 | 1.482E-02 |

| | | | | | | | |
|---|---|---|---|---|---|---|---|
| PWY-3781 | aerobic respiration I (cytochrome c) | 10141.296 | 1.472 | 0.343 | 4.294 | 1.752E-05 | 2.535E-04 |
| PWY-5088 | L-glutamate degradation VIII (to propanoate) | 201.648 | -1.844 | 0.618 | -2.986 | 2.828E-03 | 1.948E-02 |
| PWY-5180 | toluene degradation I (aerobic) (via o-cresol) | 3794.583 | 1.475 | 0.284 | 5.189 | 2.109E-07 | 4.922E-06 |
| PWY-5181 | toluene degradation III (aerobic) (via p-cresol) | 427.599 | 1.711 | 0.532 | 3.215 | 1.304E-03 | 1.073E-02 |
| PWY-5182 | toluene degradation II (aerobic) (via 4-methylcatechol) | 3794.583 | 1.475 | 0.284 | 5.189 | 2.109E-07 | 4.922E-06 |
| PWY-5198 | factor 420 biosynthesis | 2.754 | -4.721 | 1.226 | -3.852 | 1.172E-04 | 1.271E-03 |
| PWY-5266 | p-cymene degradation | 2312.737 | 1.444 | 0.370 | 3.896 | 9.767E-05 | 1.087E-03 |
| PWY-5273 | p-cumate degradation | 2312.737 | 1.444 | 0.370 | 3.896 | 9.767E-05 | 1.087E-03 |
| PWY-5415 | catechol degradation I (meta-cleavage pathway) | 1829.794 | 1.291 | 0.370 | 3.486 | 4.912E-04 | 4.576E-03 |
| PWY-5419 | catechol degradation to 2-oxopent-4-enoate II | 714.726 | 3.160 | 0.410 | 7.713 | 1.224E-14 | 1.063E-12 |
| PWY-5420 | catechol degradation II (meta-cleavage pathway) | 937.214 | 2.957 | 0.382 | 7.732 | 1.055E-14 | 1.063E-12 |
| PWY-5484 | glycolysis II (from fructose 6-phosphate) | 40147.094 | -0.178 | 0.044 | -4.018 | 5.872E-05 | 7.079E-04 |
| PWY-5507 | adenosylcobalamin biosynthesis I (early cobalt insertion) | 4640.006 | 1.915 | 0.369 | 5.185 | 2.155E-07 | 4.922E-06 |
| PWY-5647 | 2-nitrobenzoate degradation I | 246.527 | 3.704 | 0.474 | 7.807 | 5.862E-15 | 1.057E-12 |
| PWY-5651 | L-tryptophan degradation to 2-amino-3-carboxymuconate semialdehyde | 328.831 | 3.754 | 0.483 | 7.779 | 7.308E-15 | 1.057E-12 |
| PWY-5654 | 2-amino-3-carboxymuconate semialdehyde degradation to 2-oxopentenoate | 296.221 | 3.122 | 0.448 | 6.971 | 3.140E-12 | 1.703E-10 |
| PWY-5659 | GDP-mannose biosynthesis | 38965.581 | -0.166 | 0.062 | -2.683 | 7.295E-03 | 4.029E-02 |
| PWY-5695 | urate biosynthesis/inosine 5'-phosphate degradation | 50771.204 | -0.095 | 0.037 | -2.605 | 9.191E-03 | 4.806E-02 |
| PWY-5823 | superpathway of CDP-glucose-derived O-antigen building blocks biosynthesis | 166.483 | -1.632 | 0.562 | -2.906 | 3.657E-03 | 2.405E-02 |
| PWY-6071 | superpathway of phenylethylamine degradation | 733.055 | 1.855 | 0.552 | 3.362 | 7.732E-04 | 6.848E-03 |
| PWY-6107 | chlorosalicylate degradation | 61.228 | 3.917 | 0.500 | 7.830 | 4.894E-15 | 1.057E-12 |
| PWY-6123 | inosine-5'-phosphate biosynthesis I | 45852.669 | -0.111 | 0.043 | -2.594 | 9.500E-03 | 4.908E-02 |
| PWY-6145 | superpathway of sialic acids and CMP-sialic acids biosynthesis | 2.478 | -4.887 | 1.865 | -2.620 | 8.801E-03 | 4.715E-02 |
| PWY-6147 | 6-hydroxymethyl-dihydropterin diphosphate biosynthesis I | 34107.142 | -0.219 | 0.073 | -2.997 | 2.730E-03 | 1.911E-02 |
| PWY-6167 | flavin biosynthesis II (archaea) | 11.160 | -9.583 | 1.805 | -5.310 | 1.098E-07 | 3.178E-06 |
| PWY-6174 | mevalonate pathway II (archaea) | 3.427 | -4.929 | 1.051 | -4.688 | 2.759E-06 | 5.443E-05 |
| PWY-6185 | 4-methylcatechol degradation (ortho cleavage) | 283.925 | 1.667 | 0.543 | 3.070 | 2.142E-03 | 1.549E-02 |
| PWY-6339 | syringate degradation | 412.943 | 3.189 | 0.560 | 5.696 | 1.226E-08 | 4.094E-07 |

| | | | | | | | |
|---|---|---|---|---|---|---|---|
| PWY-6397 | mycolyl-arabinogalactan-peptidoglycan complex biosynthesis | 0.355 | -2.020 | 0.534 | -3.786 | 1.532E-04 | 1.622E-03 |
| PWY-6404 | superpathway of mycolyl-arabinogalactan-peptidoglycan complex biosynthesis | 0.941 | -3.007 | 0.595 | -5.054 | 4.320E-07 | 9.374E-06 |
| PWY-6467 | Kdo transfer to lipid IVA III (Chlamydia) | 16777.842 | -0.523 | 0.184 | -2.840 | 4.507E-03 | 2.794E-02 |
| PWY-6505 | L-tryptophan degradation XII (Geobacillus) | 402.798 | 3.646 | 0.476 | 7.655 | 1.928E-14 | 1.195E-12 |
| PWY-6562 | norspermidine biosynthesis | 1212.200 | 1.321 | 0.451 | 2.931 | 3.378E-03 | 2.256E-02 |
| PWY-6590 | superpathway of Clostridium acetobutylicum acidogenic fermentation | 12954.282 | 0.493 | 0.168 | 2.932 | 3.363E-03 | 2.256E-02 |
| PWY-6608 | guanosine nucleotides degradation III | 29572.896 | 0.302 | 0.093 | 3.244 | 1.177E-03 | 1.002E-02 |
| PWY-6612 | superpathway of tetrahydrofolate biosynthesis | 34378.395 | -0.186 | 0.041 | -4.534 | 5.782E-06 | 1.091E-04 |
| PWY-6629 | superpathway of L-tryptophan biosynthesis | 2082.939 | -1.565 | 0.570 | -2.747 | 6.023E-03 | 3.533E-02 |
| PWY-6641 | superpathway of sulfolactate degradation | 725.382 | 2.570 | 0.433 | 5.936 | 2.916E-09 | 1.054E-07 |
| PWY-6654 | phosphopantothenate biosynthesis III | 4.455 | -4.981 | 1.521 | -3.275 | 1.058E-03 | 9.185E-03 |
| PWY-6690 | cinnamate and 3-hydroxycinnamate degradation to 2-oxopent-4-enoate | 228.844 | 2.278 | 0.560 | 4.066 | 4.782E-05 | 6.045E-04 |
| PWY-6700 | queuosine biosynthesis | 26862.916 | -0.323 | 0.093 | -3.483 | 4.956E-04 | 4.576E-03 |
| PWY-6703 | preQ0 biosynthesis | 22139.674 | -0.338 | 0.105 | -3.214 | 1.311E-03 | 1.073E-02 |
| PWY-6728 | methylaspartate cycle | 42.338 | -1.660 | 0.542 | -3.065 | 2.180E-03 | 1.551E-02 |
| PWY-6944 | androstenedione degradation | 5.711 | -2.453 | 0.625 | -3.927 | 8.594E-05 | 1.008E-03 |
| PWY-7003 | glycerol degradation to butanol | 28255.622 | 0.239 | 0.092 | 2.608 | 9.101E-03 | 4.806E-02 |
| PWY-7007 | methyl ketone biosynthesis | 13.891 | -3.542 | 0.674 | -5.258 | 1.456E-07 | 3.950E-06 |
| PWY-7209 | superpathway of pyrimidine ribonucleosides degradation | 96.089 | 5.981 | 1.108 | 5.399 | 6.686E-08 | 2.073E-06 |
| PWY-722 | nicotinate degradation I | 12.977 | 2.792 | 0.454 | 6.145 | 8.015E-10 | 3.162E-08 |
| PWY-7323 | superpathway of GDP-mannose-derived O-antigen building blocks biosynthesis | 27105.052 | -0.344 | 0.120 | -2.853 | 4.325E-03 | 2.760E-02 |
| PWY-7373 | superpathway of demethylmenaquinol-6 biosynthesis II | 1.518 | -2.418 | 0.842 | -2.871 | 4.097E-03 | 2.654E-02 |
| PWY-7376 | cob(II)yrinate a,c-diamide biosynthesis II (late cobalt incorporation) | 10.773 | -2.896 | 0.661 | -4.384 | 1.167E-05 | 1.746E-04 |
| PWY-7527 | L-methionine salvage cycle III | 557.896 | 1.147 | 0.407 | 2.814 | 4.887E-03 | 2.987E-02 |
| PWY-7539 | 6-hydroxymethyl-dihydropterin diphosphate biosynthesis III (Chlamydia) | 35516.097 | -0.216 | 0.068 | -3.153 | 1.619E-03 | 1.269E-02 |
| PWY-7616 | methanol oxidation to carbon dioxide | 103.706 | 2.952 | 0.450 | 6.559 | 5.434E-11 | 2.358E-09 |

| | | | | | | | |
|---|---|---|---|---|---|---|---|
| PWY0-1277 | 3-phenylpropanoate and 3-(3-hydroxyphenyl)propanoate degradation | 573.344 | 1.881 | 0.524 | 3.592 | 3.280E-04 | 3.235E-03 |
| PWY0-321 | phenylacetate degradation I (aerobic) | 764.834 | 1.898 | 0.558 | 3.399 | 6.764E-04 | 6.116E-03 |
| REDCITCYC | TCA cycle VIII (helicobacter) | 11272.754 | 0.886 | 0.196 | 4.509 | 6.527E-06 | 1.180E-04 |
| RHAMCAT-PWY | L-rhamnose degradation I | 19246.146 | -0.349 | 0.111 | -3.138 | 1.699E-03 | 1.294E-02 |
| THISYN-PWY | superpathway of thiamin diphosphate biosynthesis I | 27259.032 | -0.303 | 0.109 | -2.774 | 5.531E-03 | 3.289E-02 |

### Supplementary Figures

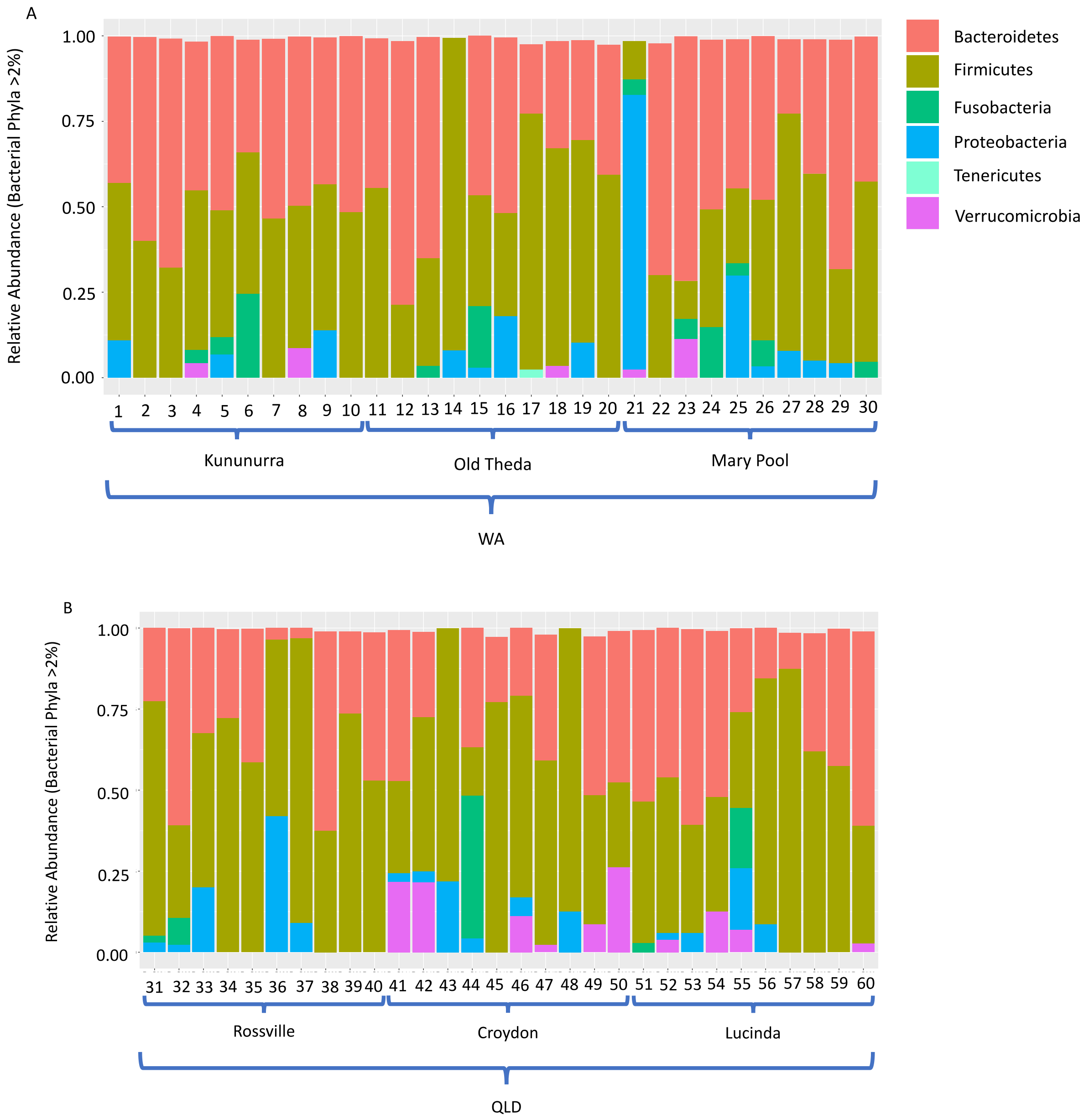


**Figure S1.** Core bacterial community composition of different individual toads. Relative abundance plot shows phyla (>2%). WA: invasion-front; QLD: range-core.

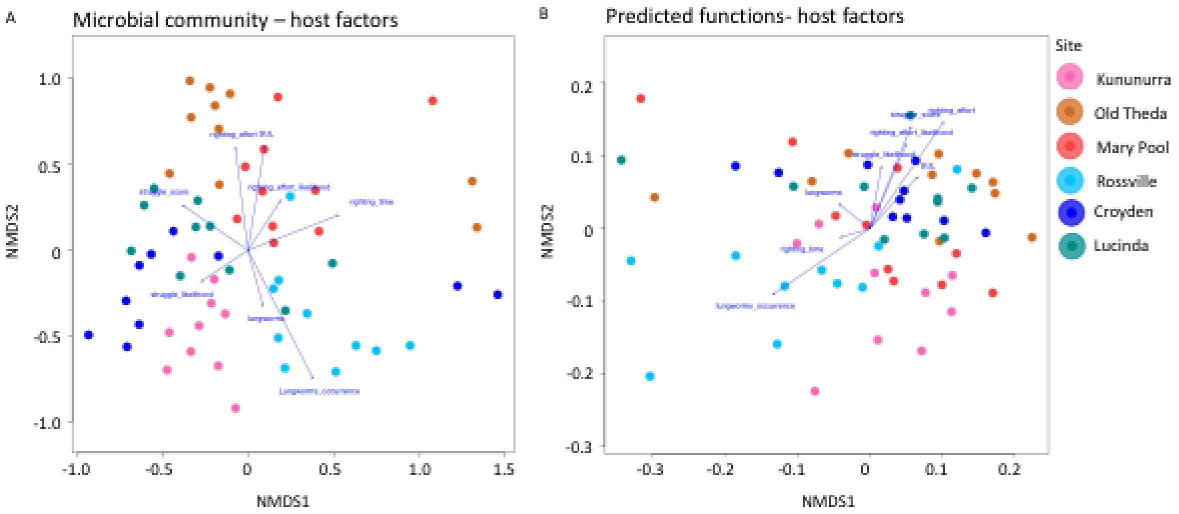


Envfit:
SUL: r2 = 0.071, p value = 0.119
Lungworms: r2 = 0.020, p value = 0.585
Occurrence of lungworms: r2 = 0.128, p value = 0.023 *
Struggle score: r2 = 0.044, p value = 0.268
Struggle likelihood: r2 = 0.021, p value = 0.555
Righting effort: r2 = 0.075, p value = 0.121
Righing effort likelihood: r2 = 0.026, p value = 0.474
Righting time: r2 = 0.062, p value = 0.174

Envfit:
SUL: r2 = 0.021, p value = 0.556
Lungworms: r2 = 0.007, p value = 0.839
Occurrence of lungworms: r2 = 0.059, p value = 0.187
Struggle score: r2 = 0.051, p value = 0.221
Struggle likelihood: r2 = 0.017, p value = 0.606
Righting effort: r2 = 0.070, p value = 0.141
Righing effort likelihood: r2 = 0.035, p value = 0.362
Righting time: r2 = 0.004, p value = 0.894

**Figure S2.** Main variables that affect the differentiation among individual cane toads' gut (A) bacterial community and (B) predicted functions. Nonmetric multidimensional scaling (nMDS) based on Bray Curtis dissimilarity in the bacterial community (stress: 0.226, A) and predicted functional profiles (stress=0.162, B). Each individual is represented by a single dot and colour coded according to sampling site (range-core: Rossville, Croyden, Lucinda; invasion-front: Kununurra, Old Theda, Mary Pool).

[8] Herlemann DP, Labrenz M, Jürgens K, Bertilsson S, Waniek JJ, Andersson AF. Transitions in

bacterial communities along the 2000 km salinity gradient of the Baltic Sea. ISME J 2011;5:1571–9. doi:10.1038/ismej.2011.41.
[9] Zhou J, Nelson TM, Rodriguez Lopez C, Sarma RR, Zhou SJ, Rollins LA. A comparison of nonlethal sampling methods for amphibian gut microbiome analyses. Mol Ecol Resour 2020;20:844–55. doi:10.1111/1755-0998.13139.
[10] Bolyen E, Rideout JR, Dillon MR, Bokulich NA, Abnet C, Al-Ghalith GA, et al. QIIME 2: Reproducible, interactive, scalable, and extensible microbiome data science 2018. doi:10.7287/peerj.preprints.27295v1.
[11] Callahan BJ, McMurdie PJ, Rosen MJ, Han AW, Johnson AJA, Holmes SP. DADA2: High-resolution sample inference from Illumina amplicon data. Nat Methods 2016;13:581–3. doi:10.1038/nmeth.3869.
[12] DeSantis TZ, Hugenholtz P, Larsen N, Rojas M, Brodie EL, Keller K, et al. Greengenes, a chimera-checked 16S rRNA gene database and workbench compatible with ARB. Appl Environ Microbiol 2006;72:5069–72. doi:10.1128/AEM.03006-05.
[13] McMurdie PJ, Holmes S. phyloseq: an R package for reproducible interactive analysis and graphics of microbiome census data. PLoS One 2013;8:e61217. doi:10.1371/journal.pone.0061217.
[14] Pielou EC. The measurement of diversity in different types of biological collections. J Theor Biol 1966;13:131–44. doi:10.1016/0022-5193(66)90013-0.
[15] Shannon CE. A mathematical theory of communication. Bell System Technical Journal 1948;27:379–423. doi:10.1002/j.1538-7305.1948.tb01338.x.
[16] Bletz MC, Goedbloed DJ, Sanchez E, Reinhardt T, Tebbe CC, Bhuju S, et al. Amphibian gut microbiota shifts differentially in community structure but converges on habitat-specific predicted functions. Nat Commun 2016;7:13699. doi:10.1038/ncomms13699.
[17] Valverde A, Makhalanyane TP, Cowan DA. Contrasting assembly processes in a bacterial metacommunity along a desiccation gradient. Front Microbiol 2014;5:668. doi:10.3389/fmicb.2014.00668.
[18] Oksanen J, Blanchet FG, Friendly M, Kindt R, Legendre P, McGlinn D, et al. vegan: Community Ecology Package. vegan: Community Ecology Package; 2019.
[19] Nath R, Pavur R. A new statistic in the one-way multivariate analysis of variance. Comput Stat Data Anal 1985;2:297–315. doi:10.1016/0167-9473(85)90003-9.
[20] Herve M. Testing and plotting procedures for biostatistics. Package "RVAideMemoire" 2018.
[21] Hochberg Y. A sharper Bonferroni procedure for multiple tests of significance. Biometrika 1988;75:800–2. doi:10.1093/biomet/75.4.800.
[22] Douglas GM, Maffei VJ, Zaneveld J, Yurgel SN, Brown JR, Taylor CM, et al. PICRUSt2: An improved and extensible approach for metagenome inference. BioRxiv 2019. doi:10.1101/672295.
[23] Love MI, Huber W, Anders S. Moderated estimation of fold change and dispersion for RNA-seq data with DESeq2. Genome Biol 2014;15:550. doi:10.1186/s13059-014-0550-8.
